# IFIH1 (MDA5) is required for innate immune detection of intron-containing RNA expressed from the HIV-1 provirus

**DOI:** 10.1101/2023.11.17.567619

**Authors:** Mehmet Hakan Guney, Karthika Nagalekshmi, Sean Matthew McCauley, Claudia Carbone, Ozkan Aydemir, Jeremy Luban

## Abstract

Antiretroviral therapy (ART) suppresses HIV-1 viremia and prevents progression to AIDS. Nonetheless, chronic inflammation is a common problem for people living with HIV-1 on ART. One possible cause of inflammation is ongoing transcription from HIV-1 proviruses, whether or not the sequences are competent for replication. Previous work has shown that intron-containing RNA expressed from the HIV-1 provirus in primary human blood cells, including CD4^+^ T cells, macrophages, and dendritic cells, activates type 1 interferon. This activation required HIV-1 *rev* and was blocked by the XPO1 (CRM1)-inhibitor leptomycin. To identify the innate immune receptor required for detection of intron-containing RNA expressed from the HIV-1 provirus, a loss-of-function screen was performed with shRNA-expressing lentivectors targeting twenty-one candidate genes in human monocyte derived dendritic cells. Among the candidate genes tested, only knockdown of XPO1 (CRM1), IFIH1 (MDA5), or MAVS prevented activation of the IFN-stimulated gene ISG15. The importance of IFIH1 protein was demonstrated by rescue of the knockdown with non-targetable IFIH1 coding sequence. Inhibition of HIV-1-induced ISG15 by the IFIH1-specific Nipah virus V protein, and by IFIH1-transdominant inhibitory CARD-deletion or phosphomimetic point mutations, indicates that IFIH1 filament formation, dephosphorylation, and association with MAVS, are all required for innate immune activation in response to HIV-1 transduction. Since both IFIH1 and DDX58 (RIG-I) signal via MAVS, the specificity of HIV-1 RNA detection by IFIH1 was demonstrated by the fact that DDX58 knockdown had no effect on activation. RNA-Seq showed that IFIH1-knockdown in dendritic cells globally disrupted the induction of IFN-stimulated genes. Finally, specific enrichment of unspliced HIV-1 RNA by IFIH1 was revealed by formaldehyde crosslinking immunoprecipitation (f-CLIP). These results demonstrate that IFIH1 is required for innate immune activation by intron-containing RNA from the HIV-1 provirus, and potentially contributes to chronic inflammation in people living with HIV-1.

## INTRODUCTION

Antiretroviral therapy (ART) suppresses HIV-1 viremia, protects CD4^+^ T cells, and prevents progression to AIDS, but it does not eliminate HIV-1 infection (1, 2). T cell activation markers that are elevated by HIV-1 infection are decreased after ART (3, 4), though systemic inflammation with activation of monocytes and macrophages remains an ongoing problem for many people living with HIV-1 on ART (5–8). Chronic immune activation is associated with non-AIDS-related complications of HIV-1 infection including elevated risk of cardiovascular disorders (9–12). In people living with HIV-1, Janus kinase inhibitor ruxolitinib decreased markers of inflammation (13), and pitavastatin reduced the risk of major adverse cardiovascular events (14). Given that inflammation is also a feature of aging, the magnitude of this public health problem is compounded by the increasing number of people living with HIV-1 who are over 50 years of age (15). Hence, better understanding of the chronic inflammation associated with HIV-1 infection would potentially aid both clinical care of individuals with, and public health responses to, this chronic disease.

Several mechanisms have been proposed to contribute to chronic immune activation in people living with HIV-1 on ART. Some inhibitors of HIV-1 protease and integrase are associated with increased rates of obesity, metabolic syndrome, and other conditions linked to inflammation (16–18). The prevalence of infection with chronic pathogens, including herpes viruses, hepatitis viruses, and tuberculosis, is higher in people living with HIV-1, and these co-infections are associated with increased immune activation and cardiovascular disease risk (19–22). Destruction of CD4^+^ T_H_17 cells during acute HIV-1 infection may weaken gut barrier function, permitting translocation of inflammatory microbial products from the intestinal lumen into the systemic circulation (23, 24). Similarly, homeostatic innate lymphoid cells within the gut lamina propria are permanently depleted by the cytokine storm that accompanies acute HIV-1 infection, contributing to disruption of gut barrier function (25, 26).

HIV-1 RNA expressed from proviruses is another factor that may contribute to chronic immune activation and inflammation in people living with HIV-1 on ART. Though ART potently suppresses HIV-1 replication, it does not eliminate proviruses established prior to initiation of ART. In people living with HIV-1 on ART, such proviruses may persist in memory CD4^+^ T cells for the life-time of the individual (1, 2, 27, 28), and possibly in other long-lived cell types such as liver macrophages (29–34). Over 95% of proviruses in people living with HIV-1 on ART have mutations which preclude production of infectious virus (35–37). Nonetheless, as much as 20% of these provirus-bearing cells express HIV-1 RNA (38–40). The amount of cell-associated HIV-1 RNA, rather than the quantity of intact HIV-1 provirus, correlates with T cell activation and plasma levels of inflammatory markers (41, 42). Collectively, these observations suggest that HIV-1 RNA promotes innate immune activation without requirement for new rounds of infection.

How the innate immune system detects HIV-1 RNA remains unclear but mechanisms may differ depending on the target cell-type and whether the infection is acute or chronic. HIV-1 RNA is detected by TLR7 during acute challenge of plasmacytoid dendritic cells (43–45), and possibly CD4^+^ T cells (46), but myeloid dendritic cells have limited expression of TLR7 and do not detect HIV-1 in the same manner (47–49). Transfection of HIV-1 genomic RNA activated RIG-I-dependent signaling (50, 51) but the relevance of RIG-I in the context of HIV-1 infection has not been confirmed. Detection of nascent HIV-1 cDNA by cyclic GMP-AMP synthase (cGAS) has been reported (52–55), while other research groups have failed to confirm the importance of cGAS for detection of HIV-1 (56–60). Such discrepancies might be explained by differences in protocol, including how HIV-1 challenge stocks are produced, the identity of the target cells, or CA amino acid residues that influence CA lattice stability (61–64).

We and others have reported that induction of innate immune signaling after HIV-1 challenge of primary human dendritic cells, macrophages, or CD4^+^ T cells requires integration, transcription from the nascent provirus, and nuclear export of intron-containing HIV-1 RNA through the Rev-CRM1 pathway (59, 60). The experiments described here had the goal of identifying the innate immune sensor of unspliced RNA produced by the HIV-1 provirus.

## RESULTS

### Loss-of-function screen in dendritic cells for genes required for detection of intron-containing RNA expressed from the HIV-1 provirus

HIV-1 transduction activates innate immune signaling and type 1 IFN in human blood CD4^+^ T cells, monocyte-derived dendritic cells, and monocyte-derived macrophages (52, 54, 55, 59, 60, 65). This host response requires HIV-1 reverse transcription, integration, transcription from the provirus, and expression of the CRM1-dependent, Rev-dependent, Rev Response Element (RRE)-containing, unspliced HIV-1 RNA (59, 60). To identify host genes required for innate immune detection of unspliced RNA expressed from the HIV-1 provirus, a targeted loss-of-function screen was performed in which twenty-one host genes encoding previously described HIV-1 RNA binding proteins (66–68), or well-characterized innate immune nucleic acid receptors, were knocked down in dendritic cells, and ISG15 induction by HIV-1 transduction was monitored by flow cytometry (59).

As previously described (59, 69), human CD14^+^ monocytes were enriched from peripheral blood mononuclear cells (PBMCs) with magnetic beads and transduced with lentivectors encoding puromycin acetyltransferase and a short hairpin RNA (shRNA) designed to knockdown transcripts from one of the candidate genes (Fig. 1A). To increase transduction efficiency, monocytes were challenged simultaneously with virus-like particles (VLPs) bearing SIV_MAC_251 Vpx (70). For the primary screen, each candidate gene was targeted by three different shRNAs, using monocytes from two blood donors. An shRNA that targets luciferase (Luc) was used as a negative control. Immediately following incubation with the knockdown vector and the Vpx^+^ VLPs, monocytes were placed in culture with IL-4 and GM-CSF produced in human cells to promote differentiation into dendritic cells. Three days after transduction, puromycin was added to the medium to select for cells that had been transduced. The dendritic cells that result from this protocol are immature and do not express ISGs (Figure 1B and references (59, 60, 65)).

**Figure 1.**
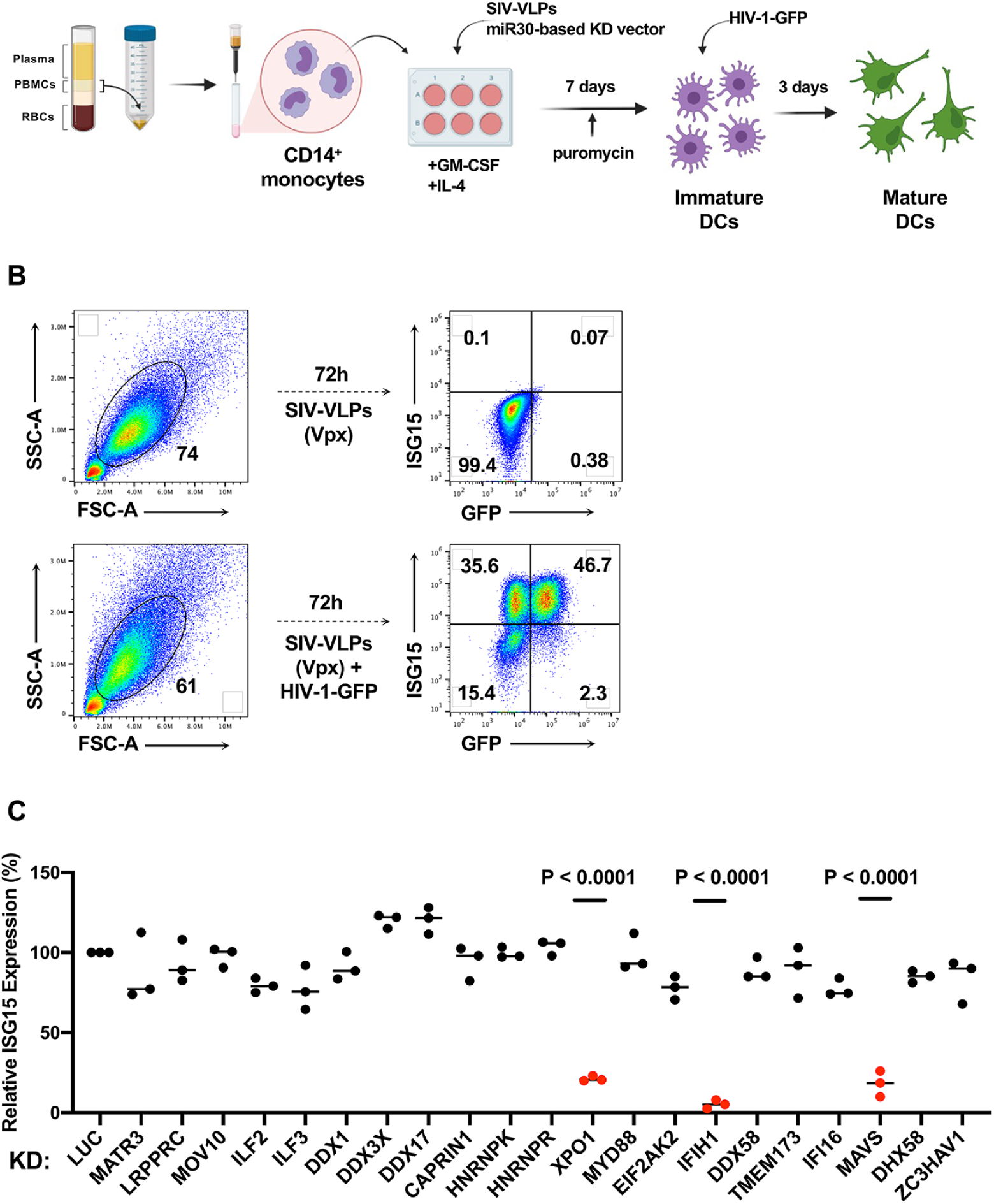
Targeted loss-of-function screen in dendritic cells identified IFIH1, MAVS, and XPO1 as required for ISG15 induction by HIV-1 transduction. **A**, Schematic showing experimental design of the knockdown screen in monocyte-derived dendritic cells. CD14^+^ monocytes transduced with shRNA-puromycin resistance vectors were differentiated into dendritic cells and selected with puromycin. Differentiated cells were transduced with HIV-1-GFP on day 7. Three days later cells were stained for intracellular ISG15 and analyzed by flow cytometry. **B**, Representative gating strategy for flow cytometry data presented in this manuscript. **C**, Loss-of-function screen in dendritic cells targeting twenty-one genes encoding HIV-1 RNA binding proteins or innate immune receptors. The plot depicts ISG15 signal upon HIV-1-GFP transduction after suppressing the indicated candidate gene relative to ISG15 signal in Luciferase control knockdown cells (n=3 shRNA target sites, n=2 donors). *P* values were determined by one-way ANOVA with Dunnett’s multiple comparisons test, relative to luciferase knockdown control.

On day seven, the resulting immature dendritic cells were challenged with single-cycle HIV-1 produced by transfection of HIV-1 proviral DNA bearing a deletion in *env* and eGFP in place of *nef* (HIV-1-GFP), along with a vesicular stomatitis virus glycoprotein (VSV G) expression plasmid (71) (Fig. 1A). VSV G pseudotyping increases transduction efficiency, though dendritic cells mature in response to transduction using the HIV-1 glycoprotein and in the absence of Vpx^+^ VLPs (59). dendritic cells were then assessed by flow cytometry for GFP and ISG15 (Fig. 1B). ISG15 was evident in both HIV-1-GFP-transduced and -untransduced dendritic cells (59), an observation explained previously as bystander activation by secreted IFN-β (65).

Of the twenty-one candidate genes targeted in our screen, only knockdown of XPO1 (Exportin 1, also called CRM1), IFIH1 (also known as MDA5), or MAVS, inhibited ISG15 induction in response to HIV-1-GFP (Fig. 1C). Reduction in ISG15 signal by knockdown of XPO1 was expected since XPO1-inhibitors block dendritic cell maturation in response to HIV-1 transduction (59, 60). Given that IFIH1 detects viral dsRNA (72, 73) and activates type 1 IFN via interaction with MAVS, IFIH1 seemed a plausible candidate for the innate immune receptor that detects unspliced HIV-1 RNA. Therefore, subsequent experiments focused on IFIH1 and MAVS. Knockdown of DDX58 (also known as RIG-I), an innate immune receptor that also signals via MAVS, had no effect on ISG15 induction by HIV-1 (Fig. 1C).

To validate the results of the primary screen, the importance of IFIH1 and MAVS for innate immune detection of HIV-1 was tested using dendritic cells from four additional blood donors. DDX58 was targeted as a negative control. Western blot of cell lysates from dendritic cells transduced with lentiviral vectors expressing 3 separate shRNAs for each gene reduced protein levels for MAVS by 6-, 4.5-, and 6-fold, for DDX58 by 10-, 14-, and 42-fold, and for IFIH1 by 8-, 22-, and 26-fold, respectively (Fig. 2A). Again, knockdown of MAVS or IFIH1 blocked the ISG15 induction that resulted from HIV-1 transduction (Fig. 2B). In contrast, DDX58 knockdown had no effect on ISG15 induction (Fig. 2B). Decreased ISG15 induction might result if shRNAs targeting MAVS or IFIH1 decreased the efficiency of subsequent HIV-1 transduction, but this was not the case (Fig. 2C).

**Figure 2.**
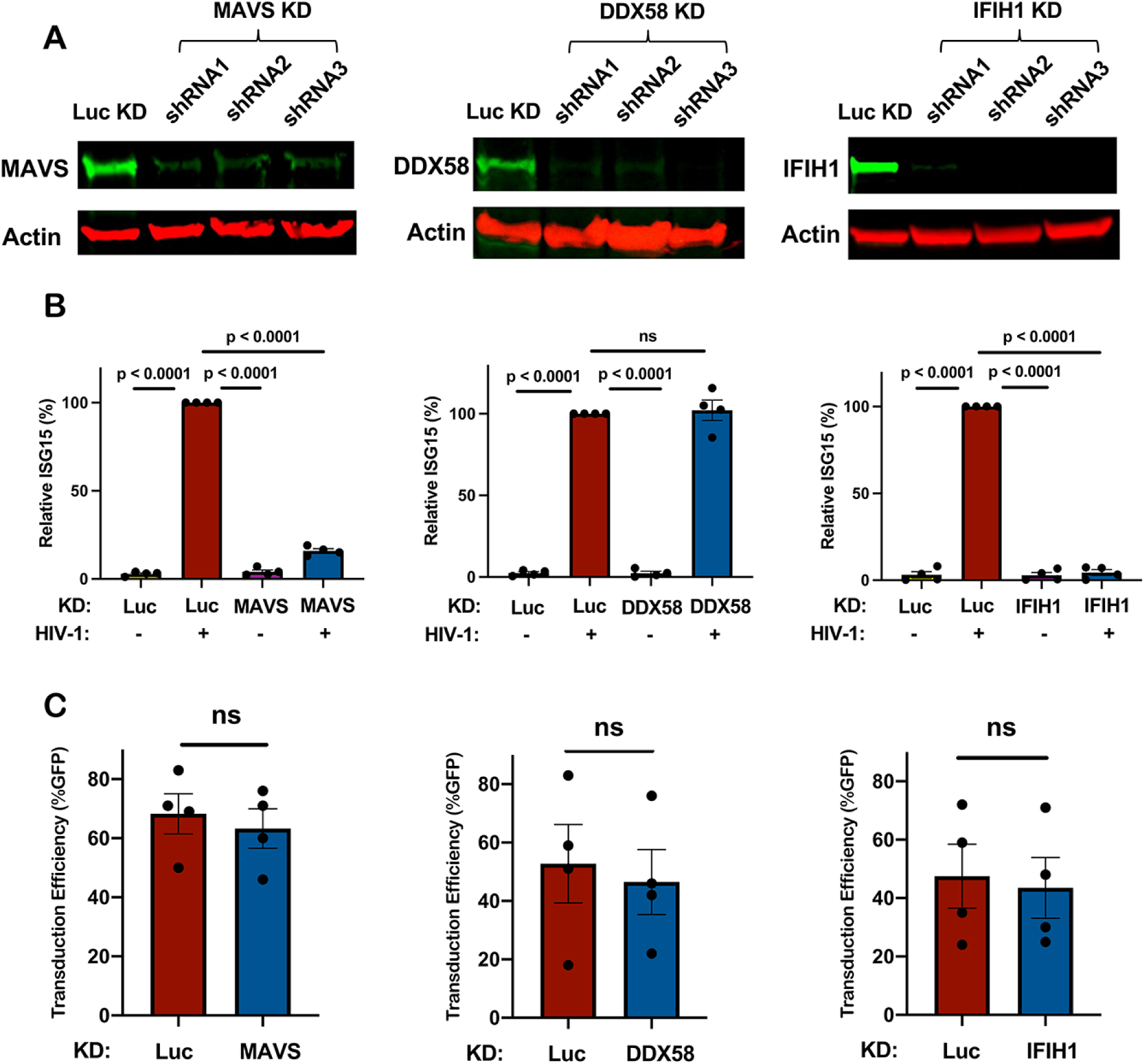
Knockdown of IFIH1 or MAVS in dendritic cells prevents ISG15 induction by HIV-1. **A**, CD14^+^ monocytes were transduced with shRNA-puromycin resistance vectors targeting MAVS, DDX58, IFIH1, or luciferase (LUC) control, as indicated, and cells were differentiated into dendritic cells under puromycin selection as in Fig. 1A. Representative immunoblots show the protein signal in the indicated samples. **B,** Differentiated cells were transduced with HIV-1-GFP for three days, and then assessed by flow cytometry for intracellular ISG15. Plots show relative ISG15 signal in the indicated knockdown cells, with and without HIV-1-GFP transduction (mean ± SEM, n=4 donors). *P* values were determined by one-way ANOVA with Tukey’s multiple comparisons test. **C,** Percentage GFP^+^ cells after HIV-1-GFP transduction of the dendritic cells in **(B)** (mean ± SEM, n=4 donors). *P* values were determined by two-tailed, paired t test.

### IFIH1 and MAVS are required to sense HIV-1 RNA in monocyte-derived macrophages

The importance of IFIH1 and MAVS for innate immune detection of HIV-1 in monocyte-derived macrophages was examined next. Macrophages were generated from CD14^+^ blood monocytes by culturing for 7 days in the presence of GM-CSF, as previously described (59, 69). Cells were transduced with lentivectors expressing shRNAs specific for IFIH1 and MAVS, and selected with puromycin, as described above for DCs. Cells were then challenged with HIV-1-GFP. As with the dendritic cells, ISG15 induction was blocked when either IFIH1 or MAVS was knocked down (Fig. 3A), and knockdown of IFIH1 or MAVS with shRNA lentivectors did not affect the efficiency of subsequent transduction by HIV-1 (Fig. 3B).

**Figure 3.**
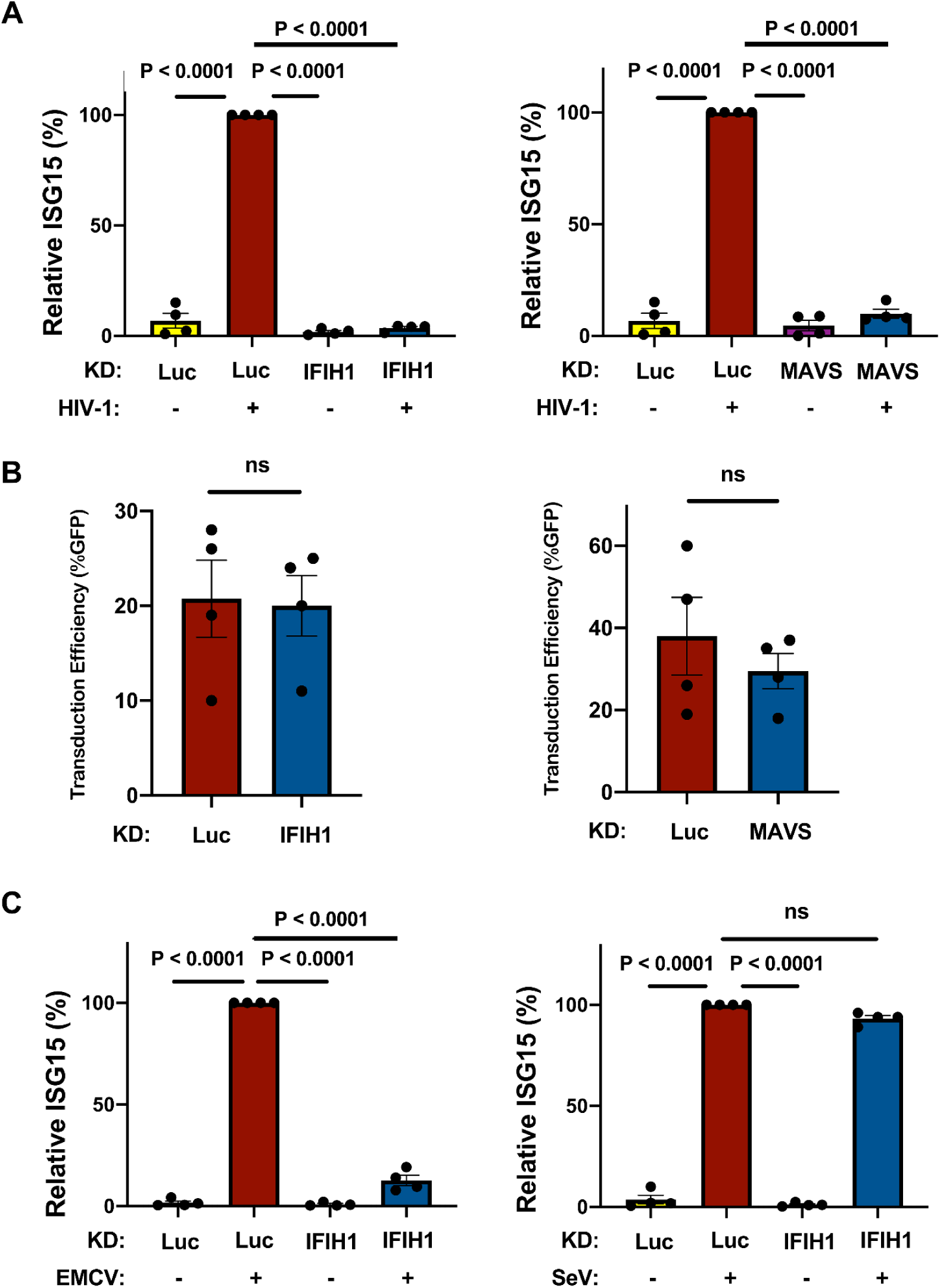
IFIH1 or MAVS knockdown prevents ISG15 induction by HIV-1 in macrophages. **A**, CD14^+^ monocytes were transduced with shRNA-puromycin resistant vectors targeting IFIH1, MAVS, or LUC control, as indicated. Cells were selected with puromycin for three days, and differentiated into macrophages in the presence of GM-CSF. Cells were transduced with HIV-1-GFP for three days and then assessed by flow cytometry for intracellular ISG15. Plots show relative ISG15 signal in the indicated knockdown cells, with and without HIV-1-GFP transduction (mean ± SEM, n=4 donors). *P* values were determined by one-way ANOVA with Tukey’s multiple comparisons test. **B,** Quantification of GFP^+^ cell populations in donors used in the experiments for **(A)** (mean ± SEM, n=4 donors). *P* values were determined by two-tailed, paired t-test. **C,** As in **(A),** except that macrophages were infected with either encephalomyocarditis virus (EMCV) or Sendai virus (SeV), instead of with HIV-1-GFP (mean ± SEM, n=4 donors). *P* values were determined by one-way ANOVA with Tukey’s multiple comparisons test.

The role of IFIH1 and DDX58 in innate immune signaling has previously been established in the context of RNA viruses other than HIV-1. More specifically, type 1 IFN induction by encephalomyocarditis virus (EMCV) is IFIH1-dependent (74), but induction by Sendai virus (SeV) is IFIH1-independent (75, 76). Challenges with these two viruses were used to determine if the behavior of the shRNA-knockdown macrophages is consistent with this literature. Indeed, EMCV and SeV each independently induced ISG15 protein production in the macrophages (Fig. 3C), and IFIH1 knockdown prevented ISG15 induction by EMCV but not by SeV (Fig. 3C). Overall, these data suggest that IFIH1 and MAVS play a critical role in innate immune sensing of nascent RNA produced by the HIV-1 provirus within human myeloid cells.

### Human IFIH1 protein is required for innate immune sensing of HIV-1 RNA in myeloid cells

To rule out potential off-target effects of IFIH1 shRNAs, and to test whether IFIH1 protein is sufficient for innate immune detection of HIV-1 RNA, a previously described all-in-one knockdown and target protein rescue vector (69) was used to simultaneously express both the shRNA that targets endogenous IFIH1 RNA, and human IFIH1 coding sequence containing silent mutations in the shRNA target sequence, from a single primary transcript. Four versions of the all-in-one vector were engineered, each containing a different combination of shRNAs, targeting either IFIH1 or Luc control, and open reading frames, encoding either non-targetable IFIH1 or heat stable antigen (HSA)/CD24 as a control open reading frame (Fig. 4A).

**Figure 4.**
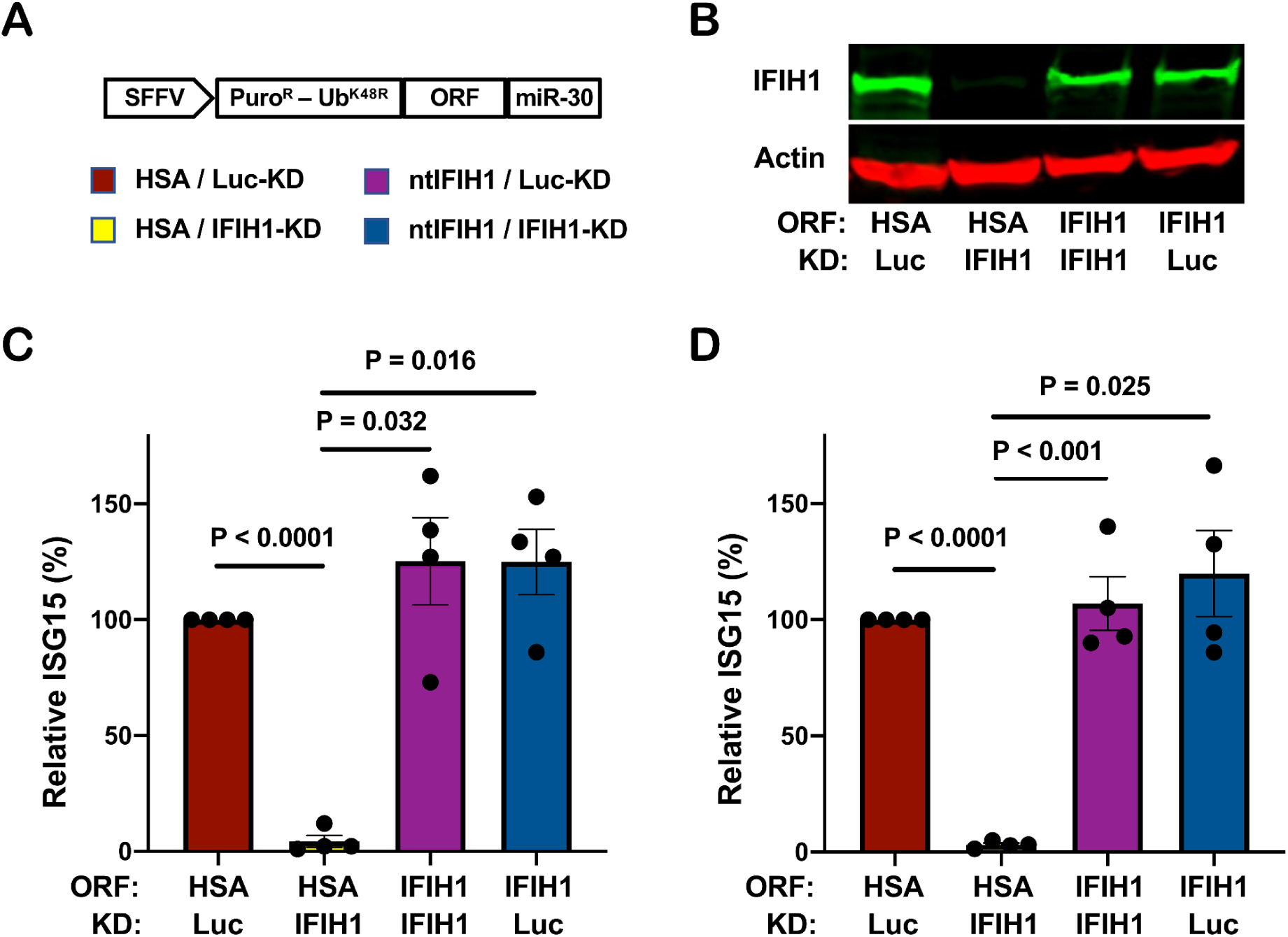
IFIH1 protein is required for ISG15 induction by HIV-1 in myeloid cells. **A**, Schematic representation of all-in-one shRNA-rescue lentivector, in which the SFFV promoter expresses a tripartite fusion of puromycin N-acetyltransferase (puroR), the K48R mutant of ubiquitin (UbK48R), and an open reading frame (ORF) encoding the gene of interest, as well as a modified miR30-based shRNA (miR-30). **B-D,** All-in-one lentivectors encoding heat stable antigen (HSA)/CD24, or non-targetable, shRNA-resistant IFIH1 coding sequence (ntIFIH1), along with shRNA targeting Luc or IFIH1 as indicated in **(A)**, were used to transduce dendritic cells **(B and C)**, or macrophages **(D)**. **B,** Representative immunoblot for quantification of IFIH1 protein levels in indicated samples. **C and D,** The percentage of ISG15^+^ cells was quantified by flow cytometry three days after HIV-1-GFP transduction, and normalized to 100% for HSA ORF/Luc KD cells (mean ± SEM, n=4 donors for each). *P* values were determined by one-way ANOVA with Tukey’s multiple comparisons test.

Dendritic cells and macrophages were transduced with each of the four versions of the all-in-one vector and selected for three days with puromycin. Cells were then challenged with the HIV-1-GFP reporter vector, and three days later assessed by western blot for steady-state levels of IFIH1 protein (Fig. 4B), and by flow cytometry for percentage of ISG15^+^ cells (Fig. 4C and D). In both dendritic cells and macrophages, transduction with the vector expressing the IFIH1-specific shRNA and the HSA coding sequence reduced IFIH1 protein levels (Fig. 4B) and abrogated ISG15 induction by HIV-1 (Fig. 4C and D). Transduction with the vector expressing the IFIH1-specific shRNA and the non-targetable IFIH1 coding sequence restored IFIH1 protein to the level in control cells (Fig. 4B) and the induction of ISG15 by HIV-1 to the control level (Fig. 4C and D). Finally, cells transduced with vector bearing shRNA targeting Luc and HSA coding sequence behaved like control cells (Fig. 4B, C, and D). These results demonstrate that IFIH1 protein is required for innate immune sensing of HIV-1 RNA in both human dendritic cells and macrophages, and that disruption of this activity by the shRNA was not due to off-target effects of the transduction vectors or the shRNAs.

### Filament formation by IFIH1 is required for innate immune detection of HIV-1

As an orthogonal method to test the specific requirement for IFIH1 in innate sensing of HIV-1, CD14^+^ monocytes were transduced with a vector engineered to express Nipah virus V protein and puromycin acetyltransferase, or with a control vector lacking the Nipah virus V coding sequence. Paramyxovirus V proteins bind to the SF2 ATP hydrolysis domain of IFIH1, but not of DDX58, and prevents the formation of filaments that is required for activation of MAVS (77–81). Transfected cells were selected with puromycin and differentiated into dendritic cells and macrophages. These puromycin-selected myeloid cell populations were then challenged with HIV-1-GFP and three days later the percentage of ISG15^+^ cells was assessed by flow cytometry.

Transduction and selection with the Nipah virus V protein-expression vector blocked ISG15-induction by HIV-1 in either dendritic cells or macrophages (Fig. 5A, B). Since IFIH1 is required for innate immune detection of EMCV (74) this virus was used as a positive control; as expected, prior transduction with the Nipah virus V-expression vector also decreased ISG15 induced by EMCV (Fig. 5C). In contrast, Nipah virus V had no effect on ISG15-induction by Sendai virus (Fig. 5D), consistent with the fact that this virus is detected by DDX58, an innate sensor that is not blocked by V protein (75, 76). Given the previously demonstrated biochemical effects of paramyxovirus V proteins (77–81), these experiments indicate that dsRNA-stimulated filament formation by IFIH1 is required for innate immune detection of HIV-1.

**Figure 5.**
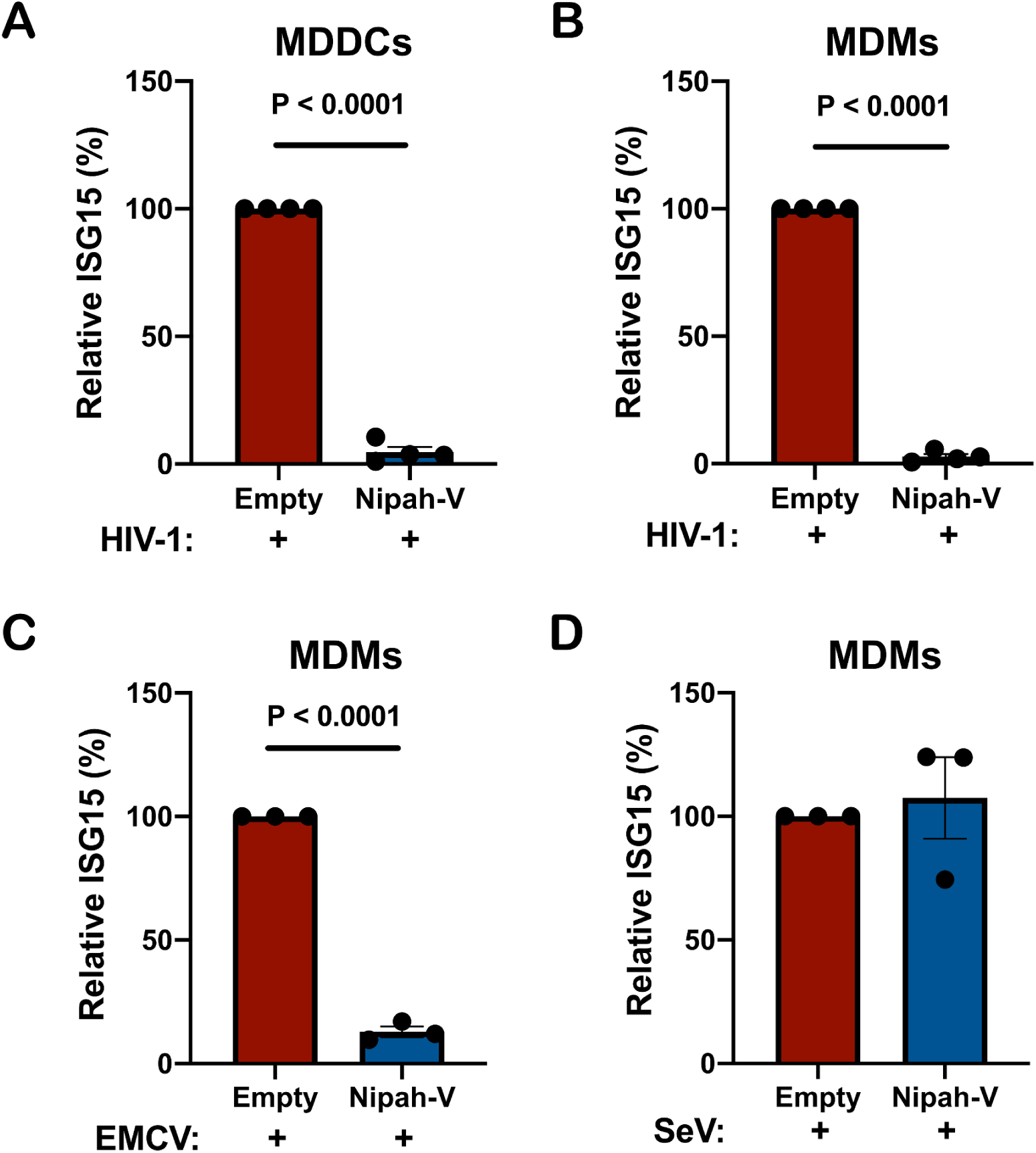
Nipah virus V protein prevents ISG15 induction by HIV-1. **A-B**, Nipah virus V protein was expressed from a lentivector in dendritic cells **(A)**, or macrophages **(B)**, and cells were subsequently transduced with HIV-1-GFP for three days. The percentage of ISG15^+^ cells was quantified by flow cytometry, and normalized to the values for cells expressing the empty vector. (mean ± SEM, n=4 donors for each). *P* values were determined by two-tailed, paired t-test. **C-D,** Nipah virus V protein was expressed in macrophages and subsequently either infected with EMCV **(C)** or Sendai virus **(D)**. The percentage of ISG15^+^ cells was measured by flow cytometry, and normalized to the values for cells expressing the empty vector (mean ± SEM, n=3 donors for each). *P* values were determined by two-tailed, paired t-test.

### IFIH1 dephosphorylation and association with MAVS is required for ISG15 activation by HIV-1

Innate immune signaling by IFIH1 requires interaction of its N-terminal tandem caspase activation and recruitment domains (CARDs) with MAVS (72, 73, 82, 83). Furthermore, phosphorylation of IFIH1 serine 88 inhibits interaction with MAVS, and phosphomimetic mutations at this position are resistant to activation (84). To test whether type 1 IFN induction by HIV-1 requires that IFIH1 signal via MAVS, an IFIH1 CARD domain deletion mutant (IFIH1Δ2CARD), and two phosphomimetic mutants, IFIH1-S88D and -S88E, were engineered into puromycin-selectable, lentiviral expression plasmids (Fig. 6A). When monocytes were transduced with a lentivector encoding WT IFIH1, or with any of the 3 IFIH1 mutants, ISG15 induction in the resulting dendritic cells was not detected in the absence of HIV-1 challenge, indicating that transduction with the IFIH1 expression vectors alone did not activate type 1 IFN (Fig. 6B). When dendritic cells were challenged with HIV-1, ISG15 induction was detected as expected in those cells that had been transduced with WT IFIH1 (Fig. 6B). In contrast, as compared to dendritic cells transduced with WT IFIH1, ISG15 induction after HIV-1 challenge was reduced 10.4-fold, 17.5-fold, and 8.9-fold in the dendritic cells that had been transduced with the IFIH1-Δ2CARD, -S88D, or -S88E mutants, respectively (Fig. 6B). Prior transduction with lentiviral vectors expressing WT IFIH1 or any of the 3 IFIH1 mutants did not change the efficiency of subsequent transduction by HIV-1 (Fig. 6C). These data demonstrate that IFIH1-dependent detection of HIV-1 requires signaling through the canonical MAVS pathway.

**Figure 6.**
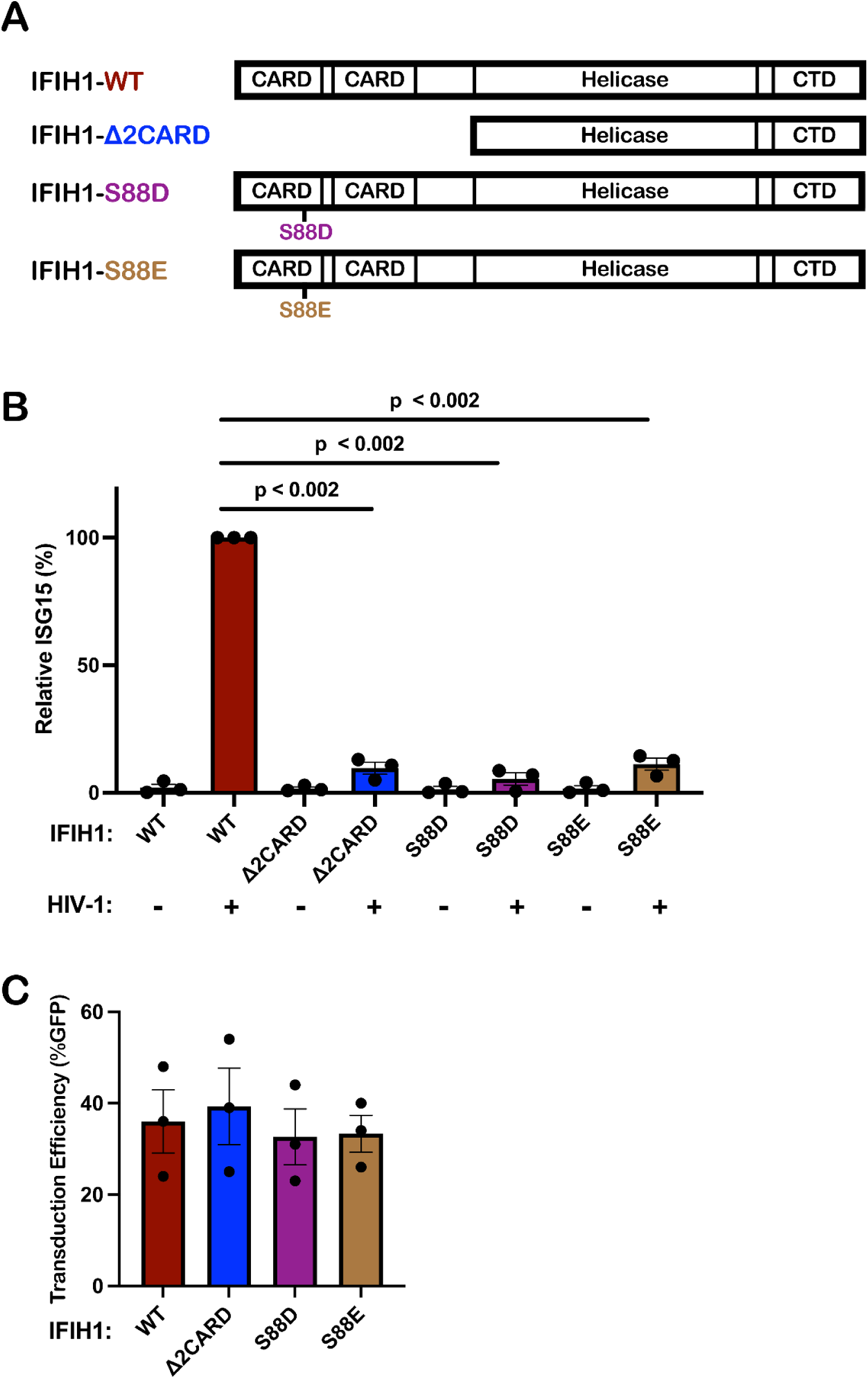
IFIH1 mutants defective for interaction with MAVS act *in trans* to abrogate ISG15 induction by HIV-1 in dendritic cells. **A**, Schematic of the IFIH1 domain structure and mutants generated here. **B,** Dendritic cells were transduced with either the vector expressing human codon optimized WT IFIH1 or one of the three indicated IFIH1 mutants. After selection with puromycin, cells were transduced with HIV-1-GFP for three days. The percentage of ISG15^+^ cells was quantified by flow cytometry, and normalized to the values for cells expressing WT IFIH1 (mean ± SEM, n=3 donors for each). *P* values were determined by one-way ANOVA with Dunnett’s multiple comparisons test. **C,** Quantification of GFP^+^ cell populations in donors used in the experiments for **(B)**, mean ± SEM, n=3 donors.

### Effect of HIV-1 transduction and IFIH1 knockdown on the dendritic cell transcriptome

The effect of HIV-1 transduction and of IFIH1 knockdown on changes to the dendritic cell transcriptome was examined next using RNA-Seq. CD14^+^ monocytes from three blood donors were each transduced with lentivectors encoding shRNAs specific for either IFIH1 or control Luc. Transduced cells were selected with puromycin and differentiated into dendritic cells. Western blot showed that, in cells expressing the IFIH1-specific shRNA from donors M27, M28, and M29, IFIH1 protein was reduced 27-, 62-, and 72-fold, respectively (Fig. 7A). The transduced dendritic cells were then challenged with HIV-1-GFP or left untreated. 48 hrs later, RNA was isolated from these cells and used to generate poly(A)-selected RNA-Seq libraries.

**Figure 7.**
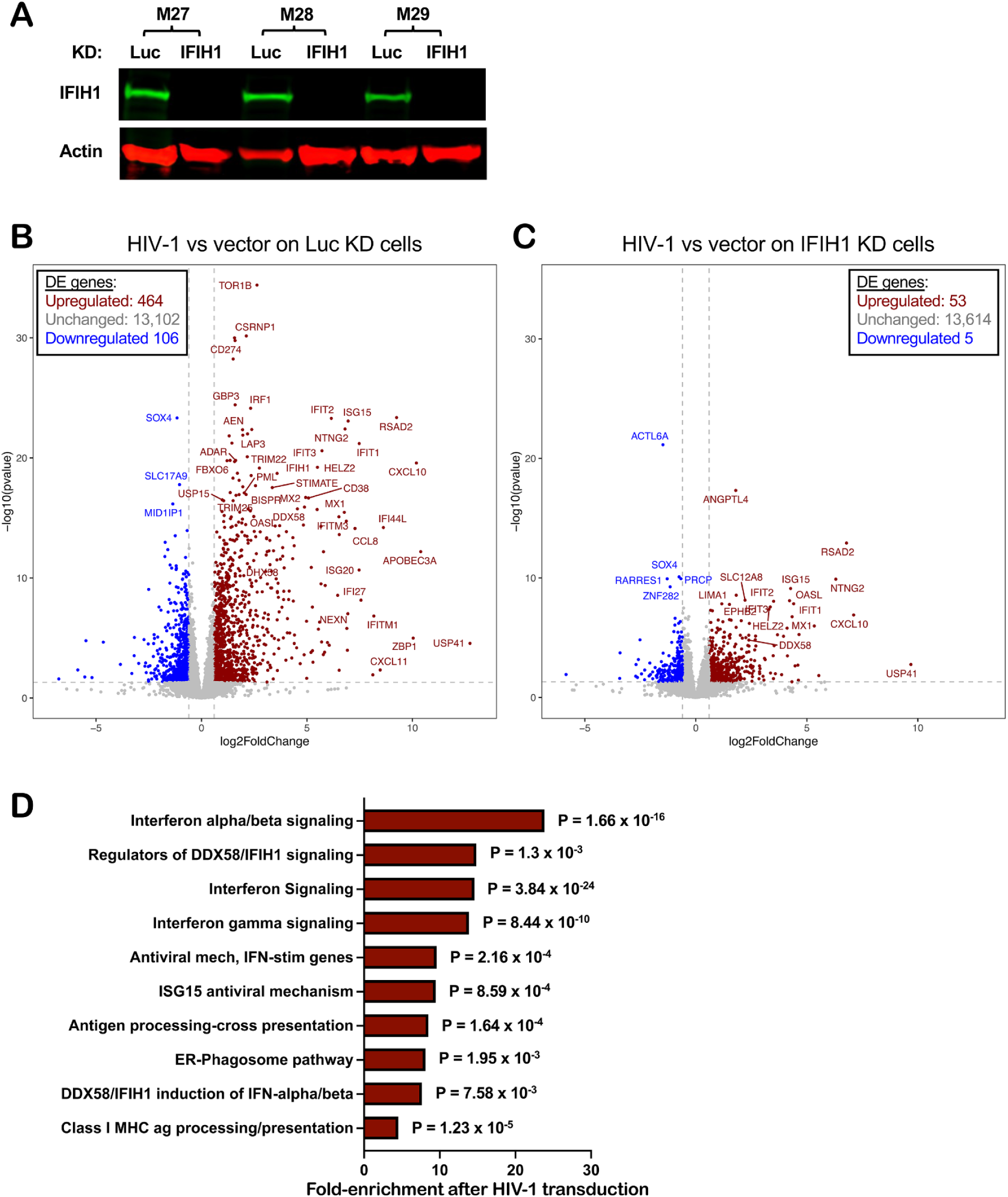
Effect of HIV-1 transduction and of IFIH1 knockdown on the dendritic cell transcriptome. CD14^+^ monocytes from three blood donors (coded M27, M28, and M29) were transduced with shRNA-puromycin resistance vectors targeting either IFIH1 or Luc control, as indicated. Cells were selected with puromycin for three days, and differentiated into dendritic cells. **A,** Immunoblot shows steady-state levels of IFIH1 protein for the indicated samples. Then, dendritic cells were either transduced with HIV-1-GFP or left untransduced. At 48 hrs, RNA was isolated from cells and poly(A)-selected libraries were generated for RNA-Seq. Volcano plot depicting differentially expressed genes after HIV-1 challenge in Luc knockdown (KD) control cells **(B),** or in IFIH1 KD cells **(C)**, as determined by RNA-Seq (log2[fold change of normalized counts]>1; P<0.01, determined by DESeq2), for the indicated samples. **D,** Reactome pathway analysis based on 276 differentially expressed genes in HIV-1 transduced Luc-KD cells in comparison to HIV-1 transduced IFIH1-KD cells. Gene Ontology-produced *P* values (as determined by Fisher’s exact test) with FDR correction (Benjamini-Hochberg method) are shown.

As compared to uninfected cells, HIV-1 transduction of control Luc knockdown cells increased expression of 464 genes and decreased expression of 106 genes (Fig. 7B, Supplementary Table 1, cut-off log_2_ ≥1 and p value ≤ 0.05). Among the upregulated interferon-stimulated genes, APOBEC3A, CXCL10, and ZBP1 were increased 10x log_2_ (p<0.0001), DDX58 was increased 4.5x log_2_ (p < 0.0001), and IFIH1 was increased 3.5x log_2_ (p < 0.0001). In contrast, the number of differentially expressed genes was greatly decreased after HIV-1 challenge of IFIH1 knockdown cells, with 53 genes increased and 5 genes decreased (Fig. 7C, Supplementary Table 2, cut-off log_2_ ≥1 and p value ≤ 0.05). Reactome pathway analysis of these differentially expressed genes showed enrichment for type 1 IFN signaling, interferon genes, and regulators of IFIH1/RIG-I signaling (Fig. 7D). Transcriptional profiling demonstrated that HIV-1 transduction of dendritic cells activates a large collection of interferon-stimulated genes, and that IFIH1 is largely required for this induction.

### Unspliced HIV-1 RNA in dendritic cells associates specifically with IFIH1

To determine if unspliced HIV-1 RNA expressed from proviruses in dendritic cells binds specifically to IFIH1, cells transduced with HIV-1-GFP were subjected to formaldehyde crosslinking immunoprecipitation (fCLIP) (85). Protein A-coated magnetic beads were incubated with affinity-purified, polyclonal rabbit anti-IFIH1 IgG, or with the equivalent anti-DDX58 IgG as a control. Beads with the bound IgG were then used to immunoprecipitate protein from formaldehyde-crosslinked dendritic cell lysates. Western blot showed that recovery of IFIH1 and DDX58 proteins was specific and quantitative, with no residual protein detected in the unbound fraction (Figure 8A). cDNA was generated from the RNA that remained bound to the beads after extensive washing, and the cDNA was then amplified by qPCR using primers specific for HIV-1 (Figure 8B). qPCR cycle threshold (Ct) values were used to calculate the fold-enrichment of HIV-1 RNA relative to the cellular gene ACTB. Unspliced HIV-1 RNA was amplified using oligonucleotide primers specific to *gag*, or to coding sequences for RT or IN (Figure 8B), and was enriched on IFIH1, 352.5-fold, 226.8-fold, and 466.7-fold, respectively (Figure 8C shows the average fold-enrichment for 3 blood donors, all p<0.0001). Spliced HIV-1 RNA was not enriched on IFIH1 (Figure 8C), whether it retained the RRE (D1A4) or not (D4A7). None of the HIV-1 RNAs were enriched on DDX58 (Figure 8C). These results demonstrate that there is specific interaction between IFIH1 and unspliced HIV-1 RNA in dendritic cells.

**Figure 8.**
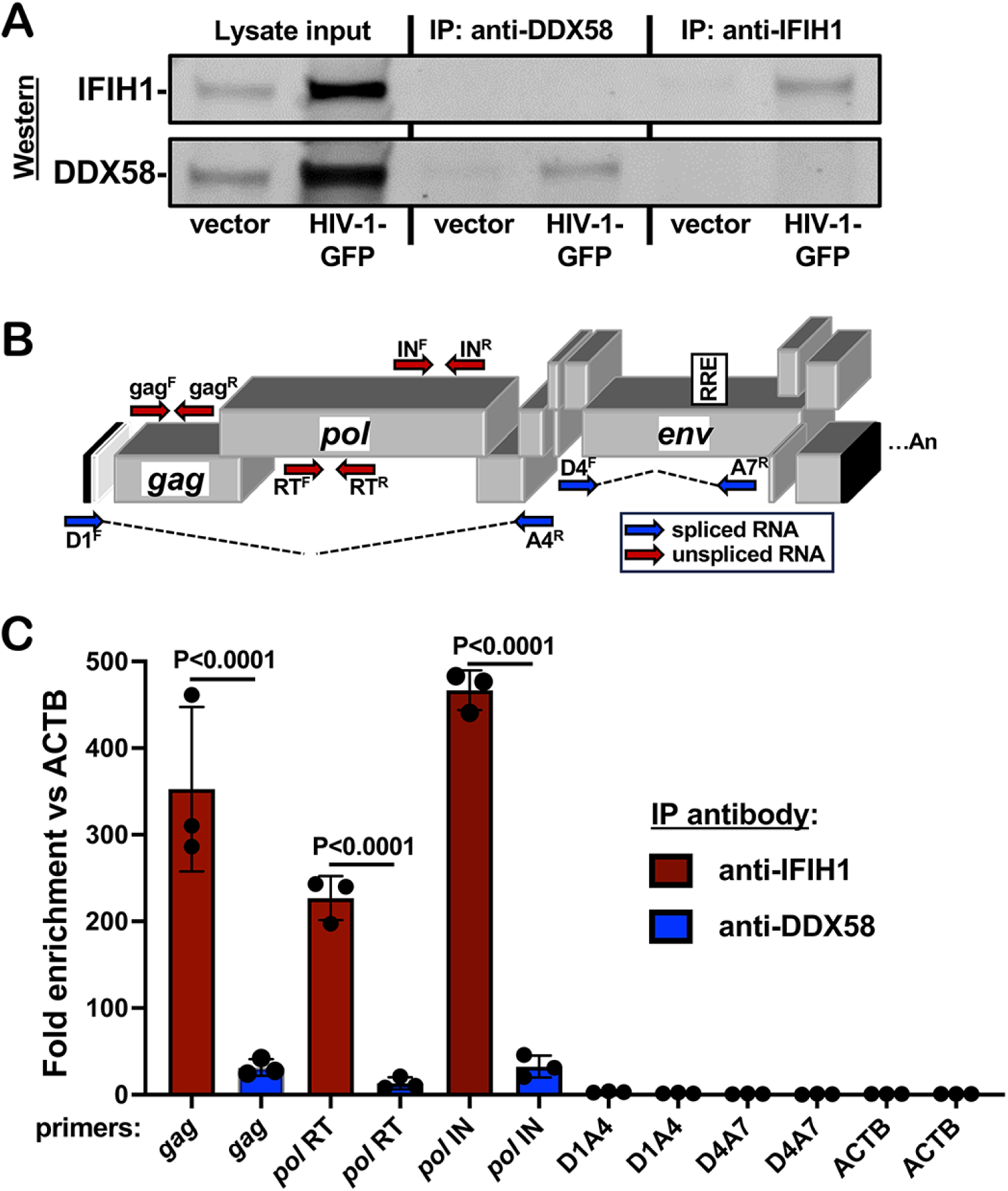
Unspliced HIV-1 RNA expressed from the provirus in dendritic cells associates specifically with IFIH1. Dendritic cells generated from three blood donors were transduced with HIV-1 GFP, or with minimal HIV-1 vector. At 72 hrs, cells were crosslinked with formaldehyde and proteins were immunoprecipitated using affinity-purified rabbit IgG specific for anti-IFIH1 (IP:anti-IFIH1) or anti-DDX58 (IP: anti-DDX58), as indicated. **A.** Western blot of cell lysate used for immunoprecipitation, anti-DDX58 immunoprecipitate, and anti-IFIH1 immunoprecipitate, probed using antibody targeting IFIH1 (upper row) or DDX58 (lower row). **B**. Schematic of the HIV-1 genome showing the location of qPCR primers that detect unspliced (red) or spliced (blue) transcripts. **C**. Fold enrichment of HIV-1 RNA in immunoprecipitates obtained with anti-IFIH1 antibody (red bars) or anti-DDX58 antibody (blue bars), as determined by RT-qPCR using primers specific for the RNAs indicated across the X axis. Each bar shows the mean value relative to ACTB for dendritic cells obtained from three individual blood donors, +/-the standard error of the mean.

## DISCUSSION

Here, a targeted loss-of-function screen based on lentiviral delivery of shRNAs in dendritic cells, identified IFIH1 and its downstream signaling molecule MAVS, as required for innate immune detection of intron-containing RNA expressed from the HIV-1 provirus (Figure 1C). These findings with IFIH1 and MAVS were confirmed using cells from multiple blood donors (Figure 2). Rescue of the knockdown with a non-targetable cDNA demonstrated the importance of IFIH1 protein for detection of HIV-1 (Figure 4), and RNA-Seq showed that IFIH1 knockdown prevented induction of the majority of interferon-stimulated genes (Figure 7). Inhibition of HIV-1-induced, interferon-stimulated genes, by the IFIH1-specific Nipah virus V protein (Figure 5), and by IFIH1-transdominant inhibitory CARD-deletion or phosphomimetic point mutations (Figure 6), indicate that IFIH1 filament formation and dephosphorylation, as well as association with MAVS, are required for innate immune activation in response to HIV-1 transduction. Though MAVS was required for innate immune detection of HIV-1 (Figure 1), and both IFIH1 and DDX58 activate type 1 interferon via MAVS (86–89), the specificity of HIV-1 RNA recognition was demonstrated by failure of efficient DDX58 knockdown to block innate immune activation by HIV-1 (Figures 1 and 2). Finally, as compared with a control cellular RNA, unspliced HIV-1 was enriched several hundred-fold by IFIH1 protein, but not by DDX58 (Figure 8). Spliced HIV-1 RNA was enriched by neither IFIH1 nor DDX58 (Figure 8). These results show that interferon-stimulated gene activation by transcription from the HIV-1 provirus requires IFIH1 recognition of *rev*-dependent, CRM1-dependent, intron-containing RNA, and IFIH1 filament formation, dephosphorylation, and association with MAVS.

It is believed that IFIH1 and DDX58 discriminate viral RNAs from cellular RNAs based on unique molecular features of the viral RNAs. DDX58, for example, binds to di-or triphosphate groups that are sometimes found at the 5’-end of viral RNAs, but not on cellular RNAs (90–97). It makes sense that HIV-1 RNA was not detected by DDX58 (Figures 1, 2, and 8) since HIV-1 RNA possesses either a 7-methylguanosine cap or a hypermethylated cap (98, 99). Stable interaction with IFIH1, instead, has been shown to require double-stranded RNA 1,000 nucleotides in length, such as the double-stranded RNA intermediates generated during viral replication (83, 100–103). HIV-1 is not known to have such long stretches of double-stranded RNA, and RNA-Seq generally does not detect antisense reads mapping to the HIV-1 provirus. HIV-1 RNA structural studies identify highly-structured regions with double-stranded character, but the longest stems are no longer than 100 nucleotides in length (104, 105). Perhaps complex stem-loop structures bearing both double-stranded and single-stranded structures like those reported for the 5’UTR and the RRE are sufficient for IFIH1 binding and activation, as others have reported with encephalomyocarditis virus (106).

Previous experiments in mice indicate that DHX58, the gene that encodes LGP2, potentiates the response of IFIH1 to poly(I:C) or to encephalomyocarditis virus (107, 108). Encephalomyocarditis virus activated human dendritic cells in an IFIH1-dependent manner (Figure 3C), and DHX58 expression was upregulated by HIV-1 transduction in dendritic cells (Supplementary Table 1), but DHX58 knockdown did not decrease innate immune activation of human dendritic cells in response to HIV-1 (Figure 1C).

Knockdown of XPO1, the gene encoding the nuclear export protein CRM1, blocked innate immune activation of dendritic cells by HIV-1 to a comparable extent as did knockdown of IFIH1 or MAVS (Figure 1C). This result confirmed the validity of the loss-of-function screen performed here, in that orthologous experiments reported inhibition with the CRM1-inhibitors leptomycin B and KPT-330, or with a null mutation in HIV-1 *rev* (59, 60). Rev binds to the highly structured RRE present in unspliced and singly-spliced HIV-1 transcripts, and possesses a leucine-rich peptide that binds to XPO1, interactions that are both required for nuclear export of the spliced and singly-spliced HIV-1 RNAs (109–112). Since unspliced HIV-1 RNA matures dendritic cells in an IFIH1-dependent manner, and requires Rev, the RRE, and XPO1 for nuclear export, and since IFIH1 localizes to the cytoplasm (113), it makes sense that all these factors are required for dendritic cell maturation by HIV-1. That being said, cytoplasmic localization of unspliced HIV-1 RNA is not sufficient for dendritic cell maturation; HIV-1 bearing disruptive mutations in *rev* and RRE does not activate innate immune signaling, even when nuclear export and p24 translation were fully rescued by heterologous NXF1-dependent nuclear export elements (59, 114). Additionally, the RRE is not sufficient to activate signaling since singly-spliced HIV-1 RNA that retains the RRE did not associate with IFIH1 in dendritic cells (Figure 8, D1A4). Taken together these findings suggest that innate immune detection of HIV-1 requires either HIV-1 sequences within the *gag-pol* intron between D1 and A4, or XPO1-dependent trafficking to a particular subcellular location.

While the data presented here was obtained in dendritic cells and in macrophages, previous studies with the same experimental tools and protocols demonstrated innate immune activation in response to intron-containing HIV-1 RNA in primary CD4^+^ T cells (59). The latter studies tracked upregulation of HLA-DR and the interferon-stimulated genes MX1 and IFIT1. Additionally, while transduction levels here were boosted by pseudotyping with VSV G in the presence of Vpx, previous studies showed that innate immune activation occurred after spreading infection with wild-type HIV-1 in the absence of Vpx (59). Taken together, the results here suggest that the chronic inflammation observed In people living with HIV-1 on ART may be caused by IFIH1-dependent activation of innate immune signaling in response to unspliced RNA from HIV-1 proviruses in CD4^+^ T cells and macrophages. This hypothesis is further supported by correlations between the quantity of cell-associated HIV-1 RNA and plasma levels of inflammatory markers (41, 42), and suggests that drugs which inhibit HIV-1 transcription (115) may decrease inflammation and cardiovascular disease risk in people living with HIV-1.

## METHODS

### Plasmids

Plasmids used in this study are listed with their corresponding Addgene accession numbers in Supplementary Table 3. The plasmids themselves, along with their complete nucleotide sequences, are available at https://www.addgene.org/Jeremy_Luban/.

### Human blood

Leukopaks were obtained from anonymous, healthy blood donors (New York Biologics). These experiments were reviewed by the University of Massachusetts Medical School Institutional Review Board and determined to be non-human subjects research, as per National Institutes of Health (NIH) guidelines, (http://grants.nih.gov/grants/policy/hs/faqs_aps_definitions.htm).

### Cell culture

All cells were cultured in humidified, 5% CO_2_ incubators at 37°C, and monitored for mycoplasma contamination using the Mycoplasma Detection kit (Lonza LT07-318). HEK293 cells (ATCC CRL-1573) were used for virus production and were cultured in Dulbecco’s Modified Eagle’s Medium (DMEM) supplemented with 10% heat-inactivated fetal bovine serum (FBS), 20 mM GlutaMAX (ThermoFisher), 1mM sodium pyruvate (ThermoFisher), 25 mM HEPES pH 7.2 (SigmaAldrich), and 1x MEM non-essential amino acids (ThermoFisher). Peripheral blood mononuclear cells (PBMCs) were isolated from leukopaks by gradient centrifugation on Lymphoprep (Axis-Shield Poc AS, catalogue no. AXS-1114546). To generate dendritic cells or macrophages, CD14^+^ mononuclear cells were enriched by positive selection using anti-CD14 antibody microbeads (Miltenyi, catalogue no. 130-050-201). Enriched CD14^+^ cells were plated in RPMI-1640, supplemented with 5% heat-inactivated human AB^+^ serum (Omega Scientific), 20 mM GlutaMAX, 1 mM sodium pyruvate, 1x MEM non-essential amino acids and 25 mM HEPES pH 7.2 (RPMI−HS complete), at a density of 2 x 10^6^ cells/ml for dendritic cells or 10^6^ cells/ml for macrophages. To differentiate CD14^+^ cells into dendritic cells, 1:100 human granulocyte-macrophage colony stimulating factor (hGM-CSF) and 1:100 human interleukin-4 (hIL-4)-conditioned media was added. To differentiate CD14^+^ cells into macrophages, 1:100 hGM-CSF-conditioned media was added. hGM-CSF and hIL-4 were produced from HEK293 cells stably transduced with pAIP-hGMCSF-co (Addgene no. 74168) or pAIP-hIL4-co (Addgene no. 74169), as previously described (59).

### HIV-1 vector production

24 hours before transfection, 6 x 10^5^ HEK-293E cells were plated per well in six-well plates. Total of 2.49 μg plasmid DNA with 6.25 μL TransIT LT1 transfection reagent (Mirus) in 250 μL Opti-MEM (Gibco) were used in all transfections. For two-part, single-cycle vector, 2.18 μg of pUC57mini NL4-3-Δenv-eGFP (NIH AIDS Reagent Program Cat #13906) was cotransfected with 0.31 μg pMD2.G VSV-G plasmid. Three-part, single-cycle vectors were produced by cotransfecting 1.25 μg minimal lentivector genome plasmid, 0.93 μg *gag*-*pol* expression plasmid (psPAX2), and 0.31 μg pMD2.G VSV G plasmid. Vpx-containing simian immunodeficiency virus (SIV)-VLPs were produced by transfection of 2.18 μg pSIV3+ and 0.31 μg pMD2.G plasmids. 16 hrs post-transfection, the culture media was changed to the media specific for the cells to be transduced. Viral supernatant was harvested at 72 hrs, filtered through a 0.45 µm filter, and stored at −80°C.

### Exogenous reverse transcriptase assay

Virions in the transfection supernatant were quantified by a PCR-based assay for reverse transcriptase activity (116). 5 μl transfection supernatant were lysed in 5 μL 0.25% Triton X-100, 50 mM KCl, 100 mM Tris-HCl pH 7.4, and 0.4 U/μl RNase inhibitor (RiboLock, ThermoFisher). Viral lysate was then diluted 1:100 in a buffer of 5 mM (NH_4_)_2_SO_4_, 20 mM KCl, and 20 mM Tris–HCl pH 8.3. 10 μL was then added to a single-step, RT PCR assay with 35 nM MS2 RNA (IDT) as template, 500 nM of each primer (5’-TCCTGCTCAACTTCCTGTCGAG-3’ and 5’-CACAGGTCAAACCTCCTAGGAATG-3’), and hot-start Taq (Promega) in a buffer of 20 mM Tris-Cl pH 8.3, 5 mM (NH_4_)_2_SO_4_, 20 mM KCl, 5 mM MgCl_2_, 0.1 mg/ml BSA, 1/20,000 SYBR Green I (Invitrogen), and 200 μM dNTPs. The RT-PCR reaction was carried out in a Biorad CFX96 real-time PCR detection system with the following parameters: 42°C 20 min, 95°C 2 min, and 40 cycles [95°C for 5 s, 60°C 5 s, 72°C for 15 s and acquisition at 80°C for 5 s]. 2 part vector transfections typically yielded 10^7^ RT units/µL, and 3 part vector transfections yielded 10^6^ RT units/µL.

### Transduction

For dendritic cells, 2 × 10^6^ CD14^+^ monocytes/mL were transduced with 1:4 volume of SIV-VLPs and 1:4 volume of knockdown lentivector, in RPMI. For macrophages, 10^6^ CD14^+^ monocytes/mL were transduced with 1:6 volume of SIV-VLPs and 1:6 volume of knockdown lentivector. The Vpx-containing SIV-VLPs were added to these cultures to overcome a SAMHD1 block to lentiviral transduction (117, 118). Transduced cells were selected with 3 μg/mL puromycin (InvivoGen, San Diego, CA, catalogue #ant-pr-1) for 3 days, starting 3 days post-transduction.

### Infectivity assay using single cycle viruses

For dendritic cells, 2.5 × 10^5^ cells were seeded per well, in a 48-well plate, on the day of virus challenge. Media containing VSV G-pseudotyped lentiviral vector expressing GFP (HIV-1-GFP) was added to challenge cells in a total volume of 250 μL. For macrophages, 2.5 × 10^5^ cells were seeded per well in a 24-well plate, and challenged with HIV-1-GFP in a total volume of 500 μL. 1:50 volume of SIV VLPs was also added to the medium during virus challenge of dendritic cells or macrophages. For both cell types, 10^8^ RT units/ml of viral vector was used to challenge cells. At 72 hours post-challenge, cells were harvested for flow cytometry by scraping. Cells were pelleted at 500 × g for 5 min, and fixed in a 1:4 dilution of BD Cytofix Fixation Buffer with phosphate-buffered saline (PBS) without Ca2+ and Mg2+, supplemented with 2% FBS and 0.1% NaN3.

### Non-HIV-1 virus challenges

Sendai Virus Cantell Strain was purchased from Charles River Laboratories. Infections were performed with 200 HA units/ml on dendritic cells and macrophages for 72 hours before assay by flow cytometry. EMCV was purchased from ATCC (VR-129B). Infections were performed with MOI 0.1 on dendritic cells and macrophages for 48 hours before assay by flow cytometry.

### Flow cytometry

10^5^ cells were pelleted at 500 x g for 5 min, and fixed in 100 μL BD Biosciences Cytofix/Cytoperm solution (catalogue #554714) for 20 minutes at 4°C. Cells were washed twice in 250 μL 1X BD Biosciences Perm/Wash buffer. Human ISG15 APC-conjugated antibody (R&D Sytems, IC8044A) was used at a dilution of 1:100 for intracellular staining. Data was collected on an Accuri C6 (BD Biosciences, San Jose, CA) and plotted with FlowJo software.

### Western Blot

Cells were lysed in Hypotonic Lysis Buffer: 20 mM Tris-HCl, pH 7.5, 150 mM NaCl, 10 mM EDTA, 0.5% NP-40, 0.1% Triton X-100, and complete mini protease inhibitor (Sigma-Aldrich) for 20 min on ice. The lysates were mixed 1:1 with 2 × Laemmli buffer containing 1:20-diluted 2-mercaptoethanol, boiled for 10 min, and centrifuged at 16,000 x g for 5 min at 4°C. Samples were run on 4–20% SDS-PAGE and transferred to nitrocellulose membranes. Membrane blocking, as well as antibody binding were in TBS Odyssey Blocking Buffer (Li-Cor, Lincoln, NE). Primary antibodies used were rabbit anti-IFIH1/MDA5 (1:1000 dilution; Proteintech, #21775-1-AP), rabbit anti-DDX58/RIG-I (1:1000 dilution; Cell Signaling Technology, #3743S), rabbit anti-MAVS (1:1000 dilution; Thermo Fisher, #PA-5-17256), and mouse anti-β-actin (1:1,000 dilution; Abcam, #ab3280). Goat anti-mouse-680 (Li-Cor, catalogue #925–68070) and goat anti-rabbit-800 (Li-Cor, catalogue #925–32211) as secondary antibodies were used at 1:10,000 dilutions. Blots were scanned on the Li-Cor Odyssey CLx.

### RNA Extraction and RNA-Seq

Total RNA was isolated from 10^6^ dendritic cells using RNeasy Plus Mini kit (Qiagen) with Turbo DNase (ThermoFisher) treatment between washes. 500 ng RNA from each sample was submitted to Genewiz for Standard poly(A) RNA-Seq.

### RNA-Seq Processing and Analysis

Quantification of human gene expression was performed on the DolphinNext platform (119) using RSEM (v1.3.1) software’s rsem-calculate-expression command, utilizing Star aligner (v2.6.1) and human genome version hg38 (Gencode v34 transcript set). A hybrid genome which included the HIV-1-GFP vector (pUC57mini_NLBN_dEnv_GFP) sequence as well as hg38 was used for viral gene quantification. Alignment (bam) files were converted to tdf format using Igvtools (v2.5.3) and visualized using IGV (v 2.10.0). DESeq2 (v1.30.1) was used for differential gene expression analysis. Donor IDs were included in the design formula (“∼ donor + condition”) to account for the differences between individuals. DESeq2’s rlog variance stabilization transformation was applied to the gene counts and the donor effect was removed using limma (v3.46.0) software removeBatchEffect function prior to generating the heatmap and PCA plots.

### Gene ontology analysis

Reactome pathway analysis was performed on https://reactome.org/ based on the 276 highest differentially expressed genes, based on Log2-fold change, comparing HIV-1 transduced Luc-KD cells in comparison to HIV-1 transduced IFIH1-KD cells. Gene Ontology-produced P values (as determined by Fisher’s exact test) with FDR correction (Benjamini-Hochberg method) are shown.

### Formaldehyde crosslinking immunoprecipitation

Protein A-coated magnetic Dynabeads (Thermo, 10001D) were washed 3 times with crosslinking immunoprecipitation (CLIP) buffer, Phosphate Buffered Saline (pH 7.4), 0.1% (w/v) SDS, 0.5% (w/v) deoxycholate, 0.5% (v/v) NP-40, protease inhibitor (Pierce, A32955), and RNAse inhibitor (Thermo, EO0382), and incubated with anti-MDA5 (IFIH1) antibody (Proteintech, 21775-1-AP) or anti-RIG-I (DDX58) antibody (Proteintech, 25068-1-AP) for 1 hr at 4°C. Dendritic cells were seeded at 10^6^ per mL in RPMI-HS complete, with Vpx+ SIV-VLP transfection supernatant added at a dilution of 1:4. After 1 hr, the cells were incubated with 1:4 volume of VSV G-pseudotyped, pUC57mini NL4-3-Δenv-eGFP (NIH AIDS Reagent Program Cat #13906). 72 hrs post transduction, cells were harvested and washed with ice cold PBS. Cells were crosslinked with freshly prepared 0.1% formaldehyde for 10 mins at room temperature and quenched by adding 1.5M glycine in PBS to a final concentration of 150 mM, and incubating for 10 mins at room temperature with gentle shaking. Crosslinked cells were washed with PBS, resuspended in CLIP buffer and sonicated with the Diagenode Bioruptor (10 cycles of 30 secs On, 30 secs Off, at 4°C). The lysate was centrifuged at 10,000 x g for 10 mins at 4°C. The supernatant was mixed with the antibody-coated Dynabeads, incubated on a rotator overnight at 4°C. The flowthrough was collected for western blot analysis to check for depletion of the protein of interest. The beads were washed three times with CLIP buffer. 10 µL of the resuspended beads in CLIP buffer was collected for western blot analysis to check the recovery of the protein of interest. The washed beads were incubated in proteinase K buffer (400 mM Tris-HCl (pH 7.4), 200 mM NaCl, 40 mM EDTA, 4% (w/v) SDS) with 10 µL of proteinase K (Thermo, EO0491) for 1 hr at 65°C with gentle agitation. 500 µL RNAzol was added to the supernatant and RNA was isolated by isopropanol precipitation. The precipitated RNA was washed with 75% ethanol, air dried, and resuspended in 20 µL nuclease free water. RNA was treated with DNAse I (Invitrogen, AM1907). Equal amounts of input RNA, as measured by NanoDrop 2000 (Thermo), were used to make cDNA with iScript Advanced cDNA Synthesis Kit (BioRad, 1725037). cDNA was amplified by qPCR with iTaq Universal SYBR Green Supermix (BioRad, 1725121), using HIV-1-specific primers (*gag*, *pol* RT, *pol* IN, D1A4, and D4A7) or primers targeting the cellular gene ACTB (primer sequences in Supplementary Table 3), with a BioRad CFX96 Real Time PCR head on a C1000 Touch Thermocycler, with the following settings: 95°C for 30 s, then {95°C for 10 s, 60°C for 30 s} x 40 cycles, followed by a melt curve from 65 to 95° C with increments of 0.5° C.

## Statistical Analysis

Experimental n values and information regarding specific statistical tests can be found in the figure legends. All statistical analyses were performed using PRISM 8.2 software (GraphPad Software, La Jolla, CA).

## Data availability

The data that support the findings of this study are available within the manuscript and in its supplementary information data files. RNA-Seq datasets generated here can be found at the NCBI Gene Expression Omnibus (GEO): GSE201250. The plasmids described in Table 1, along with their complete nucleotide sequences, are available at https://www.addgene.org/Jeremy_Luban/.

## ACKNOWLEDGEMENTS

We are grateful to the many anonymous blood donors who contributed leukocytes that were used in this project. This work was supported by NIH Grant R37AI147868 and U54AI170856 to J.L.

