## Supplemental Tables 1 to 4 for "IFIH1 (MDA5) is required for innate immune detection of intron-containing RNA expressed from the HIV-1 provirus"

**Supplementary Table 1. Differentially expressed genes in dendritic cells with luciferase control knockdown: effect of HIV-1-GFP vs minimal HIV-1 vector**

| Gene | log2FoldChange | p value | adj p value |
|---|---|---|---|
| **USP41** | 12.73 | 2.78E-05 | 4.36E-04 |
| **APOBEC3A** | 10.39 | 6.43E-13 | 6.98E-11 |
| **CXCL10** | 10.19 | 2.67E-20 | 1.37E-17 |
| **ZBP1** | 10.04 | 1.02E-05 | 1.85E-04 |
| **RSAD2** | 9.25 | 4.36E-24 | 6.84E-21 |
| **IFI44L** | 8.62 | 6.29E-15 | 1.11E-12 |
| **CXCL11** | 8.47 | 4.59E-03 | 3.03E-02 |
| **IFITM1** | 8.17 | 1.48E-07 | 4.59E-06 |
| **IFI27** | 7.56 | 7.10E-09 | 3.02E-07 |
| **IFIT1** | 7.48 | 6.36E-22 | 4.49E-19 |
| **ISG20** | 7.47 | 2.17E-11 | 1.71E-09 |
| **CCL8** | 7.28 | 7.50E-15 | 1.31E-12 |
| **ISG15** | 6.95 | 8.54E-24 | 1.04E-20 |
| **NEXN** | 6.94 | 9.50E-08 | 3.12E-06 |
| **FAM47E-STBD1** | 6.92 | 1.05E-04 | 1.35E-03 |
| **IFITM3** | 6.87 | 1.85E-15 | 3.94E-13 |
| **NTNG2** | 6.8 | 3.90E-23 | 4.29E-20 |
| **MX1** | 6.77 | 3.39E-16 | 8.12E-14 |
| **CMPK2** | 6.53 | 2.42E-14 | 3.87E-12 |
| **TNFSF10** | 6.51 | 8.03E-16 | 1.82E-13 |
| **HES4** | 6.46 | 2.78E-09 | 1.29E-07 |
| **IFIT2** | 6.15 | 5.10E-24 | 6.84E-21 |
| **GAREM1** | 5.97 | 2.39E-05 | 3.80E-04 |
| **GCH1** | 5.86 | 4.23E-10 | 2.44E-08 |
| **APOBEC3D** | 5.84 | 1.54E-04 | 1.88E-03 |
| **USP18** | 5.78 | 6.51E-13 | 6.98E-11 |
| **APOBEC3B** | 5.78 | 2.41E-04 | 2.73E-03 |
| **IFIT3** | 5.71 | 2.55E-21 | 1.71E-18 |
| **AXL** | 5.69 | 2.13E-05 | 3.44E-04 |

| | | | |
|---|---|---|---|
| **HSH2D** | 5.67 | 3.07E-10 | 1.87E-08 |
| **OASL** | 5.66 | 5.02E-15 | 9.35E-13 |
| **GBP4** | 5.6 | 4.75E-07 | 1.27E-05 |
| **HAPLN3** | 5.54 | 1.85E-06 | 4.18E-05 |
| **HELZ2** | 5.5 | 6.03E-20 | 2.99E-17 |
| **HERC5** | 5.48 | 1.99E-16 | 5.14E-14 |
| **AC068234.1** | 5.28 | 1.04E-03 | 9.29E-03 |
| **STAP1** | 5.26 | 1.14E-04 | 1.46E-03 |
| **OR52K3P** | 5.26 | 2.66E-08 | 9.79E-07 |
| **HESX1** | 5.21 | 1.32E-11 | 1.09E-09 |
| **TRIM31** | 5.19 | 1.59E-04 | 1.93E-03 |
| **CD38** | 5.07 | 2.19E-17 | 6.66E-15 |
| **CCL3L1** | 4.98 | 4.02E-07 | 1.10E-05 |
| **IL15RA** | 4.94 | 1.96E-17 | 6.12E-15 |
| **MX2** | 4.88 | 1.25E-16 | 3.43E-14 |
| **AC068631.3** | 4.87 | 2.50E-04 | 2.78E-03 |
| **EPSTI1** | 4.83 | 3.88E-15 | 7.64E-13 |
| **DDX58** | 4.55 | 1.76E-16 | 4.70E-14 |
| **TMEM52B** | 4.51 | 6.59E-05 | 9.11E-04 |
| **DSP** | 4.41 | 5.64E-06 | 1.11E-04 |
| **AIM2** | 4.4 | 2.84E-05 | 4.41E-04 |
| **IFI44** | 4.39 | 2.07E-13 | 2.47E-11 |
| **SIGLEC1** | 4.35 | 8.36E-09 | 3.51E-07 |
| **OAS3** | 4.33 | 4.39E-14 | 6.33E-12 |
| **BATF2** | 4.32 | 2.48E-12 | 2.36E-10 |
| **AC010226.1** | 4.28 | 4.63E-04 | 4.70E-03 |
| **OTOF** | 4.18 | 3.81E-06 | 7.94E-05 |
| **IGFBP4** | 4.16 | 1.23E-09 | 6.17E-08 |
| **TLR3** | 4.13 | 6.24E-09 | 2.72E-07 |
| **IL27** | 4.07 | 2.44E-07 | 7.25E-06 |
| **MFAP5** | 4.07 | 3.69E-06 | 7.73E-05 |
| **PLSCR1** | 4.03 | 1.38E-14 | 2.31E-12 |

| | | | |
|---|---|---|---|
| **AL357054.4** | 4 | 7.02E-04 | 6.68E-03 |
| **IFI6** | 3.92 | 1.44E-11 | 1.18E-09 |
| **CCL7** | 3.9 | 1.29E-09 | 6.44E-08 |
| **CD69** | 3.89 | 6.30E-09 | 2.73E-07 |
| **SELL** | 3.88 | 9.73E-06 | 1.77E-04 |
| **NT5C3A** | 3.85 | 1.54E-12 | 1.54E-10 |
| **OAS2** | 3.85 | 2.96E-13 | 3.51E-11 |
| **TMEM229B** | 3.81 | 2.73E-08 | 9.96E-07 |
| **HERC6** | 3.74 | 2.70E-12 | 2.55E-10 |
| **GBP1** | 3.71 | 4.44E-15 | 8.50E-13 |
| **NCF1B** | 3.68 | 6.66E-08 | 2.24E-06 |
| **IRF7** | 3.66 | 3.05E-14 | 4.65E-12 |
| **AC015911.7** | 3.66 | 7.45E-05 | 1.01E-03 |
| **IGF2BP3** | 3.65 | 1.45E-05 | 2.49E-04 |
| **USP30-AS1** | 3.65 | 4.10E-09 | 1.84E-07 |
| **GMPR** | 3.63 | 2.75E-10 | 1.68E-08 |
| **SP140** | 3.6 | 1.65E-09 | 7.90E-08 |
| **LYSMD2** | 3.6 | 1.35E-09 | 6.65E-08 |
| **IL15** | 3.59 | 3.07E-07 | 8.77E-06 |
| **IFIH1** | 3.58 | 1.95E-19 | 8.73E-17 |
| **DDX60** | 3.58 | 3.12E-12 | 2.90E-10 |
| **DDX60L** | 3.54 | 1.18E-10 | 7.89E-09 |
| **STIMATE-MUSTN** | 3.49 | 4.68E-15 | 8.84E-13 |
| **SDS** | 3.45 | 1.09E-05 | 1.94E-04 |
| **P2RY6** | 3.44 | 2.16E-04 | 2.51E-03 |
| **SAMD9L** | 3.41 | 5.40E-13 | 5.98E-11 |
| **STIMATE** | 3.36 | 2.95E-18 | 1.07E-15 |
| **CMYA5** | 3.34 | 6.70E-06 | 1.29E-04 |
| **ABTB2** | 3.33 | 6.81E-10 | 3.59E-08 |
| **SAMD9** | 3.32 | 2.78E-11 | 2.15E-09 |
| **SCIN** | 3.29 | 2.91E-03 | 2.12E-02 |
| **NAPSB** | 3.28 | 6.78E-10 | 3.59E-08 |

| | | | |
|---|---|---|---|
| **HMCN1** | 3.26 | 2.98E-03 | 2.16E-02 |
| **XAF1** | 3.25 | 7.81E-12 | 6.76E-10 |
| **EIF2AK2** | 3.24 | 7.43E-14 | 9.69E-12 |
| **IFI35** | 3.2 | 5.51E-13 | 6.05E-11 |
| **ATP10A** | 3.16 | 1.12E-07 | 3.61E-06 |
| **SERPING1** | 3.16 | 2.75E-07 | 8.06E-06 |
| **SECTM1** | 3.15 | 3.28E-06 | 7.01E-05 |
| **CYP19A1** | 3.14 | 4.88E-03 | 3.18E-02 |
| **IFITM2** | 3.12 | 3.13E-10 | 1.88E-08 |
| **PARP9** | 3.11 | 6.47E-13 | 6.98E-11 |
| **NCF1C** | 3.1 | 1.80E-06 | 4.09E-05 |
| **NCF1** | 3.09 | 1.90E-06 | 4.29E-05 |
| **RTP4** | 3.09 | 4.52E-14 | 6.44E-12 |
| **SP110** | 3.09 | 5.38E-14 | 7.60E-12 |
| **TRIM5** | 3.07 | 1.22E-12 | 1.26E-10 |
| **MVB12B** | 3.05 | 2.88E-11 | 2.20E-09 |
| **SCOC-AS1** | 3.04 | 3.73E-05 | 5.57E-04 |
| **IRF1-AS1** | 3 | 8.47E-08 | 2.80E-06 |
| **LY6E-DT** | 2.97 | 3.26E-05 | 4.97E-04 |
| **PDE4B** | 2.97 | 5.82E-06 | 1.14E-04 |
| **HIVEP2** | 2.95 | 1.98E-04 | 2.33E-03 |
| **PMAIP1** | 2.94 | 2.15E-07 | 6.43E-06 |
| **IDO1** | 2.9 | 7.02E-04 | 6.68E-03 |
| **TNF** | 2.9 | 1.32E-06 | 3.15E-05 |
| **CDKN2D** | 2.88 | 1.42E-09 | 6.95E-08 |
| **AP000873.2** | 2.8 | 1.43E-03 | 1.20E-02 |
| **RNF213-AS1** | 2.8 | 9.20E-11 | 6.26E-09 |
| **TAP1** | 2.78 | 1.44E-14 | 2.38E-12 |
| **AL157394.3** | 2.77 | 1.31E-03 | 1.12E-02 |
| **AC084871.4** | 2.76 | 1.55E-05 | 2.62E-04 |
| **TRIM22** | 2.74 | 7.00E-20 | 3.35E-17 |
| **GPBAR1** | 2.7 | 4.16E-05 | 6.13E-04 |

| | | | |
|---|---|---|---|
| **APOBEC3F** | 2.66 | 1.01E-06 | 2.46E-05 |
| **FAM225B** | 2.66 | 1.68E-03 | 1.36E-02 |
| **ITIH4** | 2.65 | 1.76E-05 | 2.93E-04 |
| **CCL5** | 2.65 | 1.47E-07 | 4.56E-06 |
| **FPR2** | 2.64 | 7.98E-06 | 1.50E-04 |
| **VAMP5** | 2.64 | 1.51E-09 | 7.36E-08 |
| **TOR1B** | 2.63 | 4.05E-35 | 5.43E-31 |
| **SMIM2-AS1** | 2.62 | 8.46E-03 | 4.83E-02 |
| **STAT1** | 2.6 | 9.51E-13 | 9.88E-11 |
| **DHX58** | 2.59 | 1.77E-10 | 1.14E-08 |
| **LAMP3** | 2.58 | 5.16E-08 | 1.79E-06 |
| **AC119674.2** | 2.58 | 7.95E-03 | 4.61E-02 |
| **LILRA5** | 2.57 | 1.19E-07 | 3.81E-06 |
| **TESPA1** | 2.57 | 3.89E-03 | 2.68E-02 |
| **PPM1K** | 2.57 | 3.64E-14 | 5.36E-12 |
| **RABGAP1L** | 2.56 | 2.06E-18 | 7.90E-16 |
| **CFB** | 2.55 | 7.62E-06 | 1.45E-04 |
| **LGALS2** | 2.54 | 3.77E-04 | 3.94E-03 |
| **KITLG** | 2.51 | 7.81E-03 | 4.54E-02 |
| **APOBEC3G** | 2.5 | 1.33E-14 | 2.26E-12 |
| **AZIN2** | 2.48 | 4.42E-04 | 4.52E-03 |
| **AC015911.10** | 2.48 | 3.30E-03 | 2.35E-02 |
| **CCL2** | 2.48 | 7.55E-16 | 1.74E-13 |
| **GBP5** | 2.47 | 3.55E-12 | 3.23E-10 |
| **AC004812.2** | 2.46 | 4.23E-05 | 6.22E-04 |
| **PLAAT4** | 2.46 | 2.96E-07 | 8.55E-06 |
| **MIR3945HG** | 2.45 | 1.34E-04 | 1.67E-03 |
| **CYP1B1-AS1** | 2.44 | 8.82E-07 | 2.19E-05 |
| **PTGIR** | 2.43 | 2.61E-04 | 2.90E-03 |
| **NAMPTP1** | 2.42 | 1.26E-06 | 3.02E-05 |
| **SLFN12L** | 2.41 | 2.68E-03 | 1.99E-02 |
| **AC006254.1** | 2.4 | 4.87E-04 | 4.91E-03 |

| | | | |
|---|---|---|---|
| **HPSE** | 2.4 | 7.26E-03 | 4.28E-02 |
| **BIRC3** | 2.39 | 5.08E-10 | 2.80E-08 |
| **AC004988.1** | 2.38 | 4.31E-23 | 4.29E-20 |
| **PNPT1** | 2.38 | 4.03E-03 | 2.75E-02 |
| **ZNF618** | 2.38 | 2.61E-10 | 1.62E-08 |
| **PLEKHN1** | 2.37 | 1.19E-04 | 1.52E-03 |
| **PARP10** | 2.36 | 4.61E-10 | 2.58E-08 |
| **EPHB2** | 2.35 | 2.94E-19 | 1.27E-16 |
| **BST2** | 2.34 | 7.38E-14 | 9.69E-12 |
| **OLR1** | 2.33 | 4.92E-08 | 1.72E-06 |
| **IRF1** | 2.33 | 7.31E-25 | 1.40E-21 |
| **CD1D** | 2.32 | 2.57E-16 | 6.51E-14 |
| **H4C14** | 2.31 | 1.34E-07 | 4.22E-06 |
| **TRIM69** | 2.3 | 1.47E-10 | 9.57E-09 |
| **NLRC5** | 2.29 | 7.84E-08 | 2.60E-06 |
| **H4C15** | 2.28 | 4.65E-04 | 4.72E-03 |
| **SLFN5** | 2.28 | 1.27E-11 | 1.06E-09 |
| **GRIN3A** | 2.27 | 1.10E-05 | 1.97E-04 |
| **APOL1** | 2.26 | 2.55E-09 | 1.19E-07 |
| **PSMB9** | 2.24 | 1.03E-08 | 4.25E-07 |
| **LRG1** | 2.24 | 1.47E-03 | 1.23E-02 |
| **PARP14** | 2.24 | 1.17E-08 | 4.77E-07 |
| **ANXA2R** | 2.23 | 2.26E-04 | 2.61E-03 |
| **MLXIPL** | 2.21 | 1.04E-02 | 5.60E-02 |
| **UBE2L6** | 2.21 | 1.27E-13 | 1.60E-11 |
| **NAMPT** | 2.21 | 1.79E-16 | 4.70E-14 |
| **APOL6** | 2.2 | 6.81E-14 | 9.32E-12 |
| **LAP3** | 2.18 | 1.00E-22 | 8.98E-20 |
| **ENDOD1** | 2.18 | 6.90E-05 | 9.45E-04 |
| **MDK** | 2.18 | 6.33E-12 | 5.65E-10 |
| **LGALS3BP** | 2.18 | 3.14E-07 | 8.89E-06 |
| **TRIM21** | 2.17 | 8.03E-21 | 5.13E-18 |

| | | | |
|---|---|---|---|
| **MT2A** | 2.17 | 5.97E-06 | 1.16E-04 |
| **CCDC7** | 2.16 | 4.33E-03 | 2.91E-02 |
| **AJ009632.2** | 2.16 | 1.04E-03 | 9.31E-03 |
| **NMI** | 2.16 | 3.37E-14 | 5.02E-12 |
| **TRANK1** | 2.15 | 3.90E-06 | 8.09E-05 |
| **AC015911.11** | 2.15 | 2.81E-05 | 4.38E-04 |
| **BRIP1** | 2.14 | 7.02E-04 | 6.68E-03 |
| **SLC18B1** | 2.14 | 2.25E-06 | 5.02E-05 |
| **ZNF442** | 2.13 | 9.53E-03 | 5.26E-02 |
| **CSRNP1** | 2.12 | 6.97E-31 | 4.47E-27 |
| **LAMA5** | 2.12 | 6.45E-04 | 6.23E-03 |
| **BISPR** | 2.12 | 1.12E-17 | 3.72E-15 |
| **FXYD6** | 2.11 | 2.96E-05 | 4.58E-04 |
| **KIAA1958** | 2.11 | 1.62E-09 | 7.83E-08 |
| **AC129507.1** | 2.1 | 7.79E-04 | 7.30E-03 |
| **IFIT5** | 2.1 | 3.59E-15 | 7.18E-13 |
| **APOL3** | 2.08 | 3.41E-09 | 1.54E-07 |
| **FIRRE** | 2.07 | 9.54E-07 | 2.34E-05 |
| **RNF213** | 2.07 | 1.63E-08 | 6.35E-07 |
| **TLR7** | 2.05 | 6.49E-04 | 6.26E-03 |
| **FPGT-TNNI3K** | 2.05 | 7.87E-03 | 4.56E-02 |
| **LINC01176** | 2.05 | 5.45E-05 | 7.73E-04 |
| **PHACTR4** | 2.05 | 2.93E-14 | 4.56E-12 |
| **C15orf48** | 2.04 | 3.52E-03 | 2.48E-02 |
| **GDF15** | 2.04 | 2.72E-08 | 9.94E-07 |
| **H4C8** | 2.03 | 4.10E-04 | 4.23E-03 |
| **PML** | 2.03 | 8.15E-18 | 2.80E-15 |
| **H2BC20P** | 2.02 | 2.02E-04 | 2.37E-03 |
| **HDX** | 2.02 | 1.90E-03 | 1.51E-02 |
| **LINC01134** | 2.01 | 4.39E-03 | 2.94E-02 |
| **STAT2** | 2.01 | 6.36E-11 | 4.50E-09 |
| **ITGB7** | 2 | 1.25E-04 | 1.58E-03 |

| | | | |
|---|---|---|---|
| **SLC22A4** | 1.99 | 1.98E-05 | 3.25E-04 |
| **TAGAP** | 1.99 | 1.33E-09 | 6.59E-08 |
| **RUBCN** | 1.99 | 6.42E-09 | 2.76E-07 |
| **TPI1P2** | 1.99 | 8.65E-03 | 4.90E-02 |
| **RIPOR2** | 1.97 | 7.43E-03 | 4.36E-02 |
| **PI4K2B** | 1.96 | 1.93E-06 | 4.34E-05 |
| **DTX3L** | 1.96 | 2.53E-18 | 9.44E-16 |
| **ACSM5** | 1.96 | 1.27E-22 | 1.06E-19 |
| **SLC39A14** | 1.96 | 3.36E-03 | 2.38E-02 |
| **TMPRSS13** | 1.95 | 1.21E-06 | 2.91E-05 |
| **AEN** | 1.95 | 4.48E-23 | 4.29E-20 |
| **TNFRSF10C** | 1.95 | 8.44E-04 | 7.76E-03 |
| **SYNJ2BP-COX16** | 1.95 | 9.50E-03 | 5.25E-02 |
| **TDRD7** | 1.94 | 1.25E-15 | 2.71E-13 |
| **TCF4** | 1.93 | 2.78E-09 | 1.29E-07 |
| **ALS2CL** | 1.92 | 3.99E-03 | 2.73E-02 |
| **NFKBIZ** | 1.92 | 4.59E-07 | 1.24E-05 |
| **HLA-F-AS1** | 1.92 | 5.12E-03 | 3.30E-02 |
| **OPTN** | 1.92 | 1.47E-14 | 2.39E-12 |
| **CMTR1** | 1.91 | 2.63E-15 | 5.35E-13 |
| **EXT1** | 1.9 | 2.15E-04 | 2.50E-03 |
| **AL357033.4** | 1.9 | 4.65E-13 | 5.24E-11 |
| **SHFL** | 1.9 | 1.47E-12 | 1.48E-10 |
| **HLA-F** | 1.9 | 6.29E-08 | 2.12E-06 |
| **MASTL** | 1.9 | 1.25E-08 | 5.04E-07 |
| **NFIX** | 1.88 | 1.03E-06 | 2.51E-05 |
| **SPAG1** | 1.88 | 1.52E-03 | 1.26E-02 |
| **H2BC21** | 1.87 | 3.20E-05 | 4.90E-04 |
| **SP140L** | 1.87 | 5.21E-13 | 5.82E-11 |
| **SLAMF7** | 1.86 | 4.79E-06 | 9.69E-05 |
| **AC040162.1** | 1.85 | 1.14E-06 | 2.75E-05 |
| **TAP2** | 1.85 | 2.66E-06 | 5.82E-05 |

| | | | |
|---|---|---|---|
| **GRAMD1B** | 1.84 | 4.50E-04 | 4.59E-03 |
| **KIAA1671** | 1.84 | 4.58E-03 | 3.03E-02 |
| **AL669918.1** | 1.83 | 6.63E-11 | 4.65E-09 |
| **SPATS2L** | 1.82 | 1.75E-12 | 1.72E-10 |
| **UBE2S** | 1.82 | 1.47E-06 | 3.46E-05 |
| **STAG3** | 1.81 | 1.49E-08 | 5.88E-07 |
| **CCL3** | 1.81 | 2.52E-11 | 1.97E-09 |
| **ENTPD5** | 1.81 | 3.30E-16 | 8.03E-14 |
| **CHROMR** | 1.8 | 1.86E-12 | 1.82E-10 |
| **CD80** | 1.8 | 6.34E-06 | 1.22E-04 |
| **PIWIL4** | 1.8 | 3.85E-07 | 1.06E-05 |
| **GVINP1** | 1.78 | 2.57E-06 | 5.67E-05 |
| **SHISA5** | 1.78 | 7.29E-19 | 2.96E-16 |
| **CNP** | 1.78 | 1.14E-17 | 3.72E-15 |
| **AC025423.4** | 1.78 | 3.95E-03 | 2.71E-02 |
| **HSD11B1** | 1.77 | 5.19E-05 | 7.48E-04 |
| **AL450326.1** | 1.76 | 1.09E-02 | 5.82E-02 |
| **TNFSF13B** | 1.76 | 2.22E-15 | 4.58E-13 |
| **MYO7B** | 1.75 | 4.22E-03 | 2.85E-02 |
| **BLZF1** | 1.75 | 1.66E-11 | 1.33E-09 |
| **OAS1** | 1.75 | 2.77E-07 | 8.10E-06 |
| **ODF3B** | 1.74 | 2.73E-06 | 5.95E-05 |
| **DYNLT1** | 1.74 | 3.24E-14 | 4.88E-12 |
| **AC022916.2** | 1.74 | 1.11E-02 | 5.89E-02 |
| **ZNF702P** | 1.73 | 1.00E-03 | 8.98E-03 |
| **LINC01358** | 1.73 | 5.15E-04 | 5.15E-03 |
| **C17orf67** | 1.73 | 7.97E-04 | 7.43E-03 |
| **H2AC19** | 1.71 | 5.78E-07 | 1.51E-05 |
| **GTPBP1** | 1.71 | 3.31E-09 | 1.51E-07 |
| **C1R** | 1.7 | 6.40E-04 | 6.19E-03 |
| **AC241585.2** | 1.7 | 1.86E-04 | 2.22E-03 |
| **SLC25A28** | 1.7 | 1.04E-09 | 5.27E-08 |

| | | | |
|---|---|---|---|
| **FBXO6** | 1.7 | 1.89E-19 | 8.73E-17 |
| **SAT1** | 1.7 | 3.21E-08 | 1.16E-06 |
| **PRDM11** | 1.7 | 4.88E-03 | 3.17E-02 |
| **PSME2** | 1.69 | 2.66E-14 | 4.20E-12 |
| **IL4I1** | 1.68 | 3.94E-04 | 4.10E-03 |
| **UBA7** | 1.67 | 3.01E-07 | 8.64E-06 |
| **PGAP1** | 1.67 | 5.55E-04 | 5.48E-03 |
| **CASZ1** | 1.67 | 4.85E-03 | 3.16E-02 |
| **C1GALT1** | 1.67 | 5.87E-14 | 8.19E-12 |
| **AP000331.1** | 1.67 | 1.78E-03 | 1.42E-02 |
| **MACC1** | 1.65 | 3.74E-05 | 5.57E-04 |
| **ZMYND15** | 1.64 | 8.92E-10 | 4.57E-08 |
| **CCL4** | 1.64 | 7.78E-04 | 7.29E-03 |
| **LY6E** | 1.64 | 4.13E-15 | 8.01E-13 |
| **KIAA1755** | 1.64 | 1.10E-02 | 5.85E-02 |
| **RTCB** | 1.64 | 1.31E-17 | 4.18E-15 |
| **JADE2** | 1.63 | 2.17E-10 | 1.38E-08 |
| **LRRC32** | 1.63 | 1.58E-04 | 1.92E-03 |
| **MYH3** | 1.62 | 7.74E-03 | 4.50E-02 |
| **ADAR** | 1.62 | 1.63E-20 | 9.52E-18 |
| **IFI16** | 1.62 | 1.54E-07 | 4.73E-06 |
| **LRRCC1** | 1.61 | 9.10E-04 | 8.27E-03 |
| **SPHK1** | 1.61 | 6.43E-09 | 2.76E-07 |
| **CARD16** | 1.6 | 9.36E-09 | 3.89E-07 |
| **ZNF107** | 1.6 | 7.17E-11 | 5.00E-09 |
| **GPR180** | 1.6 | 4.43E-11 | 3.26E-09 |
| **RAB37** | 1.6 | 5.76E-05 | 8.09E-04 |
| **GBP3** | 1.59 | 3.80E-25 | 8.49E-22 |
| **AL645933.5** | 1.59 | 4.83E-03 | 3.15E-02 |
| **CD274** | 1.59 | 1.68E-30 | 5.62E-27 |
| **ADRB2** | 1.59 | 5.64E-03 | 3.54E-02 |
| **CGAS** | 1.58 | 8.16E-13 | 8.55E-11 |

| | | | |
|---|---|---|---|
| **CBR1** | 1.57 | 8.21E-14 | 1.06E-11 |
| **SCAMP1-AS1** | 1.57 | 3.51E-06 | 7.45E-05 |
| **RNF19B** | 1.57 | 2.02E-20 | 1.08E-17 |
| **LILRB1** | 1.57 | 1.00E-30 | 4.47E-27 |
| **C4orf33** | 1.56 | 5.45E-07 | 1.43E-05 |
| **PHLDA2** | 1.56 | 3.13E-04 | 3.36E-03 |
| **TYMP** | 1.55 | 5.04E-05 | 7.28E-04 |
| **FAM122C** | 1.55 | 3.72E-04 | 3.91E-03 |
| **AC025171.2** | 1.54 | 1.34E-04 | 1.67E-03 |
| **AL353807.5** | 1.53 | 1.04E-06 | 2.52E-05 |
| **CXorf21** | 1.52 | 1.61E-08 | 6.30E-07 |
| **TENT5A** | 1.52 | 3.44E-06 | 7.31E-05 |
| **AC004241.1** | 1.52 | 1.60E-04 | 1.94E-03 |
| **PHF11** | 1.52 | 3.86E-10 | 2.24E-08 |
| **SLC1A4** | 1.51 | 2.45E-04 | 2.76E-03 |
| **AL133367.1** | 1.51 | 3.88E-03 | 2.68E-02 |
| **TLN2** | 1.51 | 5.49E-05 | 7.77E-04 |
| **MYD88** | 1.51 | 7.28E-14 | 9.69E-12 |
| **STX11** | 1.5 | 5.78E-29 | 1.55E-25 |
| **TRIM25** | 1.5 | 3.66E-17 | 1.05E-14 |
| **PIK3CD-AS1** | 1.5 | 1.04E-02 | 5.60E-02 |
| **NCOA7** | 1.49 | 3.41E-12 | 3.13E-10 |
| **ZNFX1** | 1.49 | 4.89E-19 | 2.05E-16 |
| **BAHCC1** | 1.49 | 3.50E-05 | 5.30E-04 |
| **GIMAP6** | 1.49 | 1.81E-04 | 2.16E-03 |
| **RAP2C-AS1** | 1.49 | 5.05E-03 | 3.26E-02 |
| **SMAD3** | 1.48 | 5.35E-08 | 1.85E-06 |
| **NUB1** | 1.48 | 4.42E-10 | 2.51E-08 |
| **AC004241.5** | 1.48 | 6.34E-03 | 3.88E-02 |
| **DUSP5** | 1.48 | 8.40E-12 | 7.18E-10 |
| **SCLT1** | 1.48 | 5.47E-06 | 1.08E-04 |
| **IL12RB1** | 1.47 | 1.43E-07 | 4.46E-06 |

| | | | |
|---|---|---|---|
| **TRIM56** | 1.47 | 7.05E-14 | 9.54E-12 |
| **SYNE2** | 1.47 | 6.64E-03 | 4.00E-02 |
| **BLNK** | 1.47 | 1.53E-05 | 2.59E-04 |
| **LILRB4** | 1.47 | 3.05E-12 | 2.86E-10 |
| **SUSD3** | 1.47 | 1.70E-03 | 1.38E-02 |
| **SSB** | 1.46 | 7.01E-12 | 6.14E-10 |
| **TMEM268** | 1.45 | 3.60E-06 | 7.63E-05 |
| **STAC3** | 1.45 | 5.19E-15 | 9.54E-13 |
| **AGRN** | 1.45 | 9.99E-06 | 1.82E-04 |
| **SWT1** | 1.45 | 5.40E-07 | 1.42E-05 |
| **CENPN** | 1.44 | 7.94E-04 | 7.41E-03 |
| **STARD4** | 1.44 | 5.84E-22 | 4.35E-19 |
| **C3AR1** | 1.44 | 2.65E-08 | 9.79E-07 |
| **FAM72B** | 1.43 | 2.48E-03 | 1.87E-02 |
| **GTF2IP12** | 1.42 | 8.28E-04 | 7.66E-03 |
| **ZNF546** | 1.42 | 2.01E-03 | 1.59E-02 |
| **KPNA5** | 1.42 | 3.66E-06 | 7.70E-05 |
| **PLAAT3** | 1.41 | 8.17E-16 | 1.83E-13 |
| **NRIP1** | 1.41 | 2.44E-12 | 2.34E-10 |
| **AF127936.1** | 1.4 | 1.16E-03 | 1.02E-02 |
| **BTN3A3** | 1.4 | 1.55E-08 | 6.11E-07 |
| **ST3GAL5-AS1** | 1.4 | 3.43E-03 | 2.42E-02 |
| **SAMD4A** | 1.39 | 4.81E-07 | 1.28E-05 |
| **CD40** | 1.39 | 6.07E-15 | 1.09E-12 |
| **LINC01232** | 1.39 | 7.71E-03 | 4.50E-02 |
| **MT1X** | 1.39 | 3.63E-04 | 3.83E-03 |
| **SRGAP2C** | 1.38 | 4.93E-09 | 2.18E-07 |
| **CALHM6** | 1.38 | 5.39E-03 | 3.43E-02 |
| **ANGPTL4** | 1.38 | 2.42E-11 | 1.89E-09 |
| **BACE2** | 1.38 | 1.45E-05 | 2.49E-04 |
| **PARP12** | 1.38 | 1.14E-15 | 2.50E-13 |
| **FUT4** | 1.38 | 1.95E-12 | 1.88E-10 |

| | | | |
|---|---|---|---|
| **FAM225A** | 1.37 | 1.62E-04 | 1.96E-03 |
| **RGL1** | 1.37 | 1.08E-11 | 9.03E-10 |
| **NAPA** | 1.36 | 6.24E-14 | 8.63E-12 |
| **BCL2L13** | 1.36 | 7.76E-18 | 2.74E-15 |
| **PANX1** | 1.36 | 1.68E-08 | 6.54E-07 |
| **LINC01426** | 1.36 | 6.25E-03 | 3.84E-02 |
| **WARS1** | 1.36 | 1.62E-20 | 9.52E-18 |
| **SNN** | 1.36 | 4.54E-10 | 2.55E-08 |
| **FAM72A** | 1.36 | 9.62E-05 | 1.26E-03 |
| **SRGAP2** | 1.35 | 5.28E-10 | 2.87E-08 |
| **HCP5** | 1.34 | 2.29E-05 | 3.66E-04 |
| **RORA** | 1.34 | 1.09E-02 | 5.82E-02 |
| **SKIL** | 1.34 | 2.50E-04 | 2.78E-03 |
| **CCDC88C** | 1.33 | 1.49E-03 | 1.23E-02 |
| **MRPL40** | 1.33 | 3.85E-09 | 1.73E-07 |
| **PCGF5** | 1.32 | 1.97E-11 | 1.57E-09 |
| **HYLS1** | 1.32 | 9.81E-03 | 5.37E-02 |
| **ASPHD2** | 1.32 | 1.92E-06 | 4.33E-05 |
| **RIMKLB** | 1.31 | 7.23E-09 | 3.06E-07 |
| **ZFAND4** | 1.31 | 5.47E-03 | 3.47E-02 |
| **HSPA1A** | 1.31 | 1.46E-22 | 1.15E-19 |
| **FKRP** | 1.31 | 5.81E-06 | 1.14E-04 |
| **RBCK1** | 1.31 | 9.19E-11 | 6.26E-09 |
| **AC245297.4** | 1.31 | 2.13E-03 | 1.65E-02 |
| **BRCA2** | 1.31 | 3.11E-03 | 2.24E-02 |
| **F3** | 1.3 | 4.54E-04 | 4.63E-03 |
| **HSPA1B** | 1.3 | 7.37E-11 | 5.12E-09 |
| **C1S** | 1.3 | 5.08E-03 | 3.28E-02 |
| **GIMAP2** | 1.3 | 5.42E-06 | 1.07E-04 |
| **CYSLTR1** | 1.3 | 5.69E-06 | 1.12E-04 |
| **METTL4** | 1.3 | 3.12E-06 | 6.71E-05 |
| **JUP** | 1.3 | 1.68E-04 | 2.03E-03 |

| | | | |
|---|---|---|---|
| **H2BC5** | 1.29 | 1.18E-03 | 1.03E-02 |
| **ENPP2** | 1.29 | 4.70E-04 | 4.76E-03 |
| **POLR1B** | 1.29 | 1.47E-05 | 2.51E-04 |
| **FRG1CP** | 1.29 | 2.16E-07 | 6.45E-06 |
| **CASP1** | 1.29 | 7.31E-09 | 3.08E-07 |
| **PSMB8** | 1.29 | 3.82E-10 | 2.22E-08 |
| **ORAI2** | 1.28 | 3.14E-12 | 2.90E-10 |
| **MNDA** | 1.28 | 1.26E-10 | 8.32E-09 |
| **ARHGAP27** | 1.27 | 3.52E-07 | 9.82E-06 |
| **TTC38** | 1.27 | 1.53E-09 | 7.41E-08 |
| **ARMH1** | 1.27 | 1.65E-05 | 2.76E-04 |
| **ZC3H12D** | 1.27 | 9.11E-03 | 5.10E-02 |
| **JAK3** | 1.27 | 9.86E-04 | 8.87E-03 |
| **RAD9A** | 1.27 | 1.16E-06 | 2.78E-05 |
| **ZBED1** | 1.26 | 1.49E-10 | 9.67E-09 |
| **CAMK1G** | 1.26 | 9.84E-03 | 5.38E-02 |
| **RNASEH1-AS1** | 1.26 | 7.62E-03 | 4.45E-02 |
| **IRAK2** | 1.26 | 1.08E-03 | 9.59E-03 |
| **AKAP7** | 1.26 | 1.47E-03 | 1.23E-02 |
| **TRIM26** | 1.26 | 4.28E-13 | 4.90E-11 |
| **SDSL** | 1.26 | 1.36E-10 | 8.87E-09 |
| **LINC01127** | 1.25 | 3.79E-03 | 2.63E-02 |
| **NADK** | 1.25 | 5.42E-11 | 3.87E-09 |
| **LRRC37A3** | 1.25 | 1.04E-04 | 1.35E-03 |
| **HEG1** | 1.25 | 9.97E-09 | 4.13E-07 |
| **MTMR11** | 1.24 | 2.84E-04 | 3.10E-03 |
| **SLFN12** | 1.24 | 5.28E-06 | 1.05E-04 |
| **PARP15** | 1.24 | 1.63E-03 | 1.33E-02 |
| **ARHGAP11B** | 1.24 | 5.81E-03 | 3.63E-02 |
| **GSTT2B** | 1.24 | 1.08E-02 | 5.79E-02 |
| **RASGEF1B** | 1.24 | 4.28E-10 | 2.44E-08 |
| **UNC93B1** | 1.24 | 7.68E-10 | 4.04E-08 |

| | | | |
|---|---|---|---|
| **IRF9** | 1.24 | 4.37E-11 | 3.24E-09 |
| **ZC3H12C** | 1.24 | 1.70E-07 | 5.17E-06 |
| **MOV10** | 1.22 | 8.73E-11 | 6.00E-09 |
| **SLC41A2** | 1.21 | 5.13E-04 | 5.13E-03 |
| **RARG** | 1.21 | 2.07E-03 | 1.62E-02 |
| **SCO2** | 1.21 | 3.91E-06 | 8.10E-05 |
| **AC244197.3** | 1.21 | 4.17E-03 | 2.82E-02 |
| **SCARB2** | 1.21 | 1.72E-20 | 9.58E-18 |
| **PPA1** | 1.21 | 3.25E-10 | 1.94E-08 |
| **SP100** | 1.21 | 5.28E-10 | 2.87E-08 |
| **H2AC6** | 1.2 | 4.00E-07 | 1.10E-05 |
| **CDC73** | 1.2 | 1.44E-13 | 1.77E-11 |
| **CDYL2** | 1.2 | 5.62E-04 | 5.54E-03 |
| **SLC25A23** | 1.2 | 8.91E-03 | 5.01E-02 |
| **ATF3** | 1.2 | 6.38E-11 | 4.50E-09 |
| **STAT4** | 1.19 | 1.60E-04 | 1.94E-03 |
| **AF117829.1** | 1.19 | 1.22E-03 | 1.05E-02 |
| **CHMP5** | 1.19 | 1.48E-13 | 1.81E-11 |
| **GBP2** | 1.19 | 1.33E-08 | 5.34E-07 |
| **SIGLEC8** | -1.21 | 1.40E-08 | 5.55E-07 |
| **ADAM12** | -1.21 | 7.97E-07 | 2.01E-05 |
| **LINC02035** | -1.22 | 3.26E-07 | 9.21E-06 |
| **FSCN1** | -1.23 | 8.34E-04 | 7.70E-03 |
| **CXCR2** | -1.24 | 2.68E-04 | 2.97E-03 |
| **KLF4** | -1.25 | 3.02E-14 | 4.65E-12 |
| **SSBP2** | -1.25 | 7.67E-05 | 1.03E-03 |
| **FMO5** | -1.26 | 5.51E-03 | 3.49E-02 |
| **LINS1** | -1.27 | 1.08E-08 | 4.43E-07 |
| **DIXDC1** | -1.27 | 2.28E-04 | 2.61E-03 |
| **AC010442.1** | -1.28 | 4.67E-03 | 3.08E-02 |
| **NAT8L** | -1.28 | 3.62E-03 | 2.53E-02 |
| **DPCD** | -1.3 | 2.20E-04 | 2.54E-03 |

| | | | |
|---|---|---|---|
| **KANTR** | -1.3 | 9.62E-03 | 5.30E-02 |
| **DACT1** | -1.31 | 2.84E-04 | 3.10E-03 |
| **CD28** | -1.31 | 2.36E-03 | 1.79E-02 |
| **PADI2** | -1.31 | 8.91E-03 | 5.01E-02 |
| **YPEL3** | -1.32 | 1.35E-06 | 3.21E-05 |
| **WNT5B** | -1.32 | 5.44E-08 | 1.88E-06 |
| **EIF4BP6** | -1.33 | 6.30E-03 | 3.86E-02 |
| **ORC6** | -1.33 | 3.22E-04 | 3.45E-03 |
| **BCL11A** | -1.34 | 2.51E-04 | 2.80E-03 |
| **MID1IP1** | -1.36 | 6.85E-17 | 1.91E-14 |
| **CENPP** | -1.38 | 3.30E-03 | 2.35E-02 |
| **CRB2** | -1.39 | 5.44E-05 | 7.73E-04 |
| **ST8SIA6** | -1.39 | 5.30E-06 | 1.05E-04 |
| **KRT79** | -1.4 | 3.30E-05 | 5.03E-04 |
| **CYGB** | -1.41 | 1.50E-11 | 1.23E-09 |
| **AC132812.1** | -1.43 | 4.06E-03 | 2.77E-02 |
| **DNAL1** | -1.44 | 3.76E-04 | 3.93E-03 |
| **IER3** | -1.45 | 1.20E-08 | 4.86E-07 |
| **OBSL1** | -1.46 | 2.14E-03 | 1.66E-02 |
| **NFE2** | -1.47 | 1.28E-08 | 5.15E-07 |
| **PPBP** | -1.47 | 1.32E-03 | 1.13E-02 |
| **AC126474.2** | -1.47 | 3.78E-04 | 3.95E-03 |
| **SPACA9** | -1.48 | 1.17E-03 | 1.02E-02 |
| **TMEM45B** | -1.48 | 1.48E-06 | 3.48E-05 |
| **PCARE** | -1.49 | 8.26E-12 | 7.10E-10 |
| **ENHO** | -1.5 | 2.68E-10 | 1.66E-08 |
| **LINC02610** | -1.51 | 4.10E-03 | 2.79E-02 |
| **PRKAR2B** | -1.51 | 4.19E-04 | 4.30E-03 |
| **AC009690.1** | -1.51 | 1.61E-03 | 1.32E-02 |
| **STK39** | -1.54 | 6.26E-09 | 2.73E-07 |
| **AC124068.1** | -1.57 | 6.92E-07 | 1.77E-05 |
| **APBA1** | -1.57 | 5.57E-03 | 3.51E-02 |

| | | | |
|---|---|---|---|
| **CXCR1** | -1.58 | 8.76E-07 | 2.17E-05 |
| **IL22RA2** | -1.58 | 6.88E-08 | 2.30E-06 |
| **PPM1H** | -1.59 | 5.15E-03 | 3.31E-02 |
| **IRS2** | -1.6 | 4.40E-13 | 5.00E-11 |
| **PDE2A** | -1.62 | 3.62E-04 | 3.82E-03 |
| **CHTF18** | -1.64 | 5.95E-03 | 3.70E-02 |
| **AC144831.1** | -1.65 | 2.52E-06 | 5.56E-05 |
| **CCL28** | -1.65 | 9.98E-05 | 1.30E-03 |
| **MFSD6L** | -1.65 | 2.22E-03 | 1.71E-02 |
| **SLC25A25-AS1** | -1.66 | 6.05E-03 | 3.74E-02 |
| **TDRD9** | -1.68 | 8.10E-04 | 7.54E-03 |
| **TCHH** | -1.68 | 6.91E-05 | 9.46E-04 |
| **ADGRB2** | -1.7 | 1.04E-02 | 5.60E-02 |
| **ADAMTS15** | -1.71 | 1.30E-06 | 3.10E-05 |
| **FAM126A** | -1.72 | 1.07E-13 | 1.37E-11 |
| **SESN3** | -1.74 | 1.64E-12 | 1.63E-10 |
| **ROR1-AS1** | -1.74 | 6.85E-04 | 6.57E-03 |
| **ERVFRD-1** | -1.75 | 8.94E-03 | 5.02E-02 |
| **ABLIM1** | -1.78 | 1.13E-02 | 6.00E-02 |
| **SRGAP1** | -1.79 | 8.04E-03 | 4.65E-02 |
| **ANLN** | -1.8 | 1.84E-03 | 1.47E-02 |
| **ASF1B** | -1.81 | 7.60E-04 | 7.15E-03 |
| **MYORG** | -1.86 | 5.63E-04 | 5.54E-03 |
| **AC135048.1** | -1.86 | 2.44E-04 | 2.75E-03 |
| **CENPU** | -1.87 | 1.07E-03 | 9.49E-03 |
| **CACNB4** | -1.88 | 4.87E-04 | 4.91E-03 |
| **RAB33A** | -1.88 | 2.11E-05 | 3.43E-04 |
| **GINS4** | -1.92 | 4.80E-08 | 1.68E-06 |
| **RIMBP3B** | -1.94 | 4.47E-04 | 4.57E-03 |
| **SMAD6** | -1.97 | 8.10E-05 | 1.09E-03 |
| **MBOAT2** | -1.97 | 8.50E-04 | 7.80E-03 |
| **AC096921.2** | -1.99 | 9.84E-03 | 5.38E-02 |

| | | | |
|---|---|---|---|
| **KBTBD11** | -2.01 | 2.61E-08 | 9.74E-07 |
| **PEG3** | -2.04 | 5.33E-03 | 3.39E-02 |
| **SDC1** | -2.05 | 2.08E-04 | 2.43E-03 |
| **PRORSD1P** | -2.08 | 2.11E-03 | 1.64E-02 |
| **TTC28** | -2.08 | 8.34E-06 | 1.56E-04 |
| **E2F1** | -2.1 | 1.12E-04 | 1.44E-03 |
| **KPNA2P3** | -2.16 | 3.55E-03 | 2.49E-02 |
| **ZNF835** | -2.21 | 4.40E-03 | 2.94E-02 |
| **AHNAK2** | -2.23 | 1.08E-05 | 1.93E-04 |
| **DUOX1** | -2.25 | 5.28E-06 | 1.05E-04 |
| **SLC16A10** | -2.26 | 8.62E-03 | 4.89E-02 |
| **MKI67** | -2.27 | 2.87E-05 | 4.46E-04 |
| **TFAP4** | -2.28 | 3.45E-04 | 3.66E-03 |
| **AC243960.1** | -2.39 | 8.10E-03 | 4.67E-02 |
| **LINC00921** | -2.44 | 2.02E-03 | 1.59E-02 |
| **IGSF3** | -2.5 | 8.91E-10 | 4.57E-08 |
| **ZNF571-AS1** | -2.52 | 2.38E-03 | 1.80E-02 |
| **GASK1A** | -2.53 | 3.11E-05 | 4.78E-04 |
| **ADGRG6** | -2.62 | 7.61E-03 | 4.45E-02 |
| **CSGALNACT1** | -2.65 | 1.19E-03 | 1.04E-02 |
| **NPHP1** | -2.69 | 2.22E-03 | 1.71E-02 |
| **KIAA1614** | -2.73 | 1.35E-05 | 2.36E-04 |
| **TACSTD2** | -2.98 | 1.30E-03 | 1.11E-02 |
| **TMEM154** | -3.18 | 8.96E-06 | 1.65E-04 |
| **WFDC21P** | -3.19 | 2.99E-04 | 3.23E-03 |
| **CCDC80** | -3.3 | 8.99E-03 | 5.04E-02 |
| **AP001931.2** | -3.84 | 1.57E-03 | 1.29E-02 |
| **SLX1B** | -4.66 | 2.22E-05 | 3.57E-04 |
| **SLC18A2** | -5.48 | 1.68E-05 | 2.81E-04 |
| **AC008878.3** | -5.87 | 4.56E-03 | 3.02E-02 |

**Supplementary Table 2. Differentially expressed genes in dendritic cells with IFIH1 knockdown: effect of HIV-1-GFP vs minimal HIV-1 vector**

| Gene | log2FoldChange | pvalue | padj |
|---|---|---|---|
| CXCL10 | 7.11 | 1.26E-07 | 6.93E-05 |
| RSAD2 | 6.79 | 1.21E-13 | 5.08E-10 |
| NTNG2 | 6.31 | 1.28E-10 | 2.30E-07 |
| IFI44L | 5.34 | 1.05E-06 | 3.40E-04 |
| IFIT1 | 4.41 | 1.45E-08 | 1.23E-05 |
| MX1 | 4.34 | 1.77E-07 | 8.94E-05 |
| ISG15 | 4.27 | 7.69E-10 | 1.08E-06 |
| OASL | 4.22 | 7.98E-09 | 8.38E-06 |
| ISG20 | 4.13 | 3.33E-04 | 2.22E-02 |
| CMPK2 | 4.12 | 1.59E-06 | 4.56E-04 |
| IFITM3 | 3.95 | 7.64E-06 | 1.51E-03 |
| USP18 | 3.68 | 5.69E-06 | 1.26E-03 |
| IFIT2 | 3.51 | 8.94E-09 | 8.67E-06 |
| LAMA2 | 3.49 | 3.24E-04 | 2.20E-02 |
| IFIT3 | 3.36 | 2.69E-08 | 1.99E-05 |
| HELZ2 | 3.3 | 4.24E-08 | 2.81E-05 |
| RASAL2-AS1 | 3.13 | 6.90E-04 | 3.46E-02 |
| HERC5 | 2.71 | 4.92E-05 | 5.59E-03 |
| IFI44 | 2.69 | 7.32E-06 | 1.47E-03 |
| EPSTI1 | 2.68 | 1.47E-05 | 2.37E-03 |
| TSPOAP1 | 2.66 | 1.06E-04 | 9.51E-03 |
| COL1A1 | 2.65 | 8.29E-04 | 3.91E-02 |
| ALS2CL | 2.49 | 6.91E-04 | 3.46E-02 |
| IRF7 | 2.42 | 6.41E-07 | 2.18E-04 |
| MX2 | 2.4 | 5.00E-05 | 5.63E-03 |
| BATF2 | 2.39 | 1.52E-04 | 1.27E-02 |
| DDX58 | 2.39 | 1.59E-05 | 2.42E-03 |
| OAS2 | 2.39 | 6.24E-06 | 1.34E-03 |
| OAS3 | 2.35 | 4.50E-05 | 5.30E-03 |

| | | | |
|---|---|---|---|
| **RNF144A** | 2.31 | 9.86E-04 | 4.25E-02 |
| **SP140** | 2.29 | 1.69E-04 | 1.39E-02 |
| **RAB43P1** | 2.25 | 9.91E-06 | 1.79E-03 |
| **SLC12A8** | 2.22 | 7.28E-09 | 8.35E-06 |
| **FHIT** | 2.18 | 9.24E-04 | 4.08E-02 |
| **THAP9** | 2.15 | 3.76E-04 | 2.37E-02 |
| **XAF1** | 2.1 | 1.09E-05 | 1.88E-03 |
| **CCL5** | 2.08 | 4.71E-04 | 2.75E-02 |
| **UXT-AS1** | 2.06 | 8.81E-05 | 8.17E-03 |
| **GMPR** | 2.05 | 4.69E-04 | 2.75E-02 |
| **OLR1** | 2.05 | 3.78E-07 | 1.49E-04 |
| **IFI6** | 2.04 | 4.51E-04 | 2.68E-02 |
| **SLC16A1-AS1** | 2.01 | 6.25E-06 | 1.34E-03 |
| **GSTM3** | 1.95 | 7.25E-04 | 3.60E-02 |
| **APOBEC3F** | 1.95 | 4.30E-04 | 2.63E-02 |
| **FCN1** | 1.92 | 8.87E-04 | 4.04E-02 |
| **IL15RA** | 1.88 | 9.26E-04 | 4.08E-02 |
| **ARMH1** | 1.82 | 2.79E-09 | 3.52E-06 |
| **MTMR11** | 1.82 | 1.36E-06 | 4.19E-04 |
| **PLSCR1** | 1.82 | 5.50E-04 | 2.95E-02 |
| **ABTB2** | 1.81 | 1.12E-03 | 4.50E-02 |
| **ANGPTL4** | 1.8 | 4.78E-18 | 3.01E-14 |
| **EIF2AK2** | 1.8 | 3.51E-05 | 4.34E-03 |
| **DDX60L** | 1.8 | 1.14E-03 | 4.52E-02 |
| **ACTL6A** | -1.49 | 7.21E-22 | 9.10E-18 |
| **TIMM23B-AGAP6** | -1.56 | 1.01E-03 | 4.29E-02 |
| **MSH5-SAPCD1** | -2.08 | 1.32E-03 | 5.06E-02 |
| **MBOAT2** | -2.52 | 1.51E-05 | 2.37E-03 |
| **GCOM1** | -2.59 | 6.71E-04 | 3.41E-02 |

**Supplementary Table 3. Plasmids used in this study**

| Plasmid Name | Purpose | Notes | Source |
|---|---|---|---|
| pAIP hGMCSF co | Stable 293 cell line expressing human cytokine GM-CSF | SFFV promoter expresses codon-optimized human GM-CSF and puromycin N-acetyltransferase | Addgene #74168 |
| pAIP hIL4 co | Stable 293 cell line expressing human cytokine IL4 | SFFV promoter expresses codon-optimized human IL4 and puromycin N-acetyltransferase | Addgene #74169 |
| pSIV Δpsi/Δenv/ΔVif/ΔVpr | $SIV_{MAC251}$ *gag-pol/vpx* | Production of SIV-VLPs containing Vpx protein | Addgene #132928 |
| pMD2.G | VSV Glycoprotein | Pseudotype HIV-1 vectors with VSV Glycoprotein | Addgene #12259 |
| psPAX2 | HIV-1 gag-pol | Encodes gag structural proteins and pol enzymes to generate virion particles to generate 3-part lentiviral vector | Addgene #12260 |
| pUC57mini NL4-3 Δenv eGFP | HIV-1 clade B molecular clone | Molecular clone of NL4-3 with deletion of 79 nucleotides following the Env signal peptide and eGFP in place of nef. "HIV-1-GFP" in the manuscript | NIH ARP #13906 |
| pPU HSA miR30 L1221 | All-in-one knockdown-rescue lentivector for luciferase (control) knockdown | SFFV promoter expresses puromycin N-acetyltransferase, exogenous HSA protein, and miR30-based shRNA targeting luciferase. Target sequence: 5′-CTTGTCGATGAGAGCGTTTGTA-3′; negative control in knockdown-rescue experiment | Addgene #174221 |
| pPU HSA miR30 MDA5 | All-in-one knockdown-rescue lentivector for MDA5 knockdown | SFFV promoter expresses puromycin N-acetyltransferase, exogenous HSA protein, and miR30-based shRNA targeting MDA5. Target sequence: 5′-TTTATACATCATCTTCTCTCGG-3′ | Addgene #174222 |

| | | | |
|---|---|---|---|
| pPU huMDA5 miR30 L1221 | All-in-one knockdown-rescue lentivector for huMDA5 expression and luciferase (control) knockdown | SFFV promoter expresses puromycin N-acetyltransferase, exogenous huMDA5 protein, and miR30-based shRNA targeting luciferase.<br>Target sequence:<br>5′-CTTGTCGATGAGAGCGTTTGTA-3′ | Addgene #174223 |
| pPU huMDA5 miR30 MDA5 | All-in-one knockdown-rescue lentivector for huMDA5 expression and MDA5 knockdown | SFFV promoter expresses puromycin N-acetyltransferase, exogenous huMDA5 protein, and miR30-based shRNA targeting MDA5.<br>Target sequence:<br>5′-TTTATACATCATCTTCTCTCGG-3′ | Addgene #174224 |
| pscALPSpuro MDA5Δ2CARD | Lentivector expressing human MDA5 that lacks 2 CARD domains | Encodes codon-optimized huMDA5 with a deletion of 2xCARD domains | Addgene #174225 |
| pscALPSpuro MDA5-S88D | Lentivector expressing human MDA5 S88D mutant | Encodes codon-optimized huMDA5 S88D mutant | Addgene #174226 |
| pscALPSpuro MDA5-S88E | Lentivector expressing human MDA5 S88E mutant | Encodes codon-optimized huMDA5 S88E mutant | Addgene #174227 |
| pscALPSpuro Nipah V | Lentivector expressing Nipah virus V protein | Encodes codon-optimized Nipah virus V protein | Addgene #174228 |
| pAPM miR30 L1221 | Lentivector for luciferase (control) knockdown | SFFV promoter expresses puromycin N-acetyltransferase and miR30-based shRNA targeting luciferase. Target sequence: 5′-TACAAACGCTCTCATCGACAAG-3′; negative control for knockdown | Addgene #115846 |
| pAPM miR30 CAPRIN1-ts1 | Lentivector for CAPRIN1 knockdown | SFFV promoter expresses puromycin N-acetyltransferase and miR30-based shRNA targeting CAPRIN1. Target sequence: 5′-TCTAGATTCAGAATGAACTGGA-3′ | Addgene #174229 |

| | | | |
|---|---|---|---|
| pAPM miR30 CAPRIN1-ts2 | Lentivector for CAPRIN1 knockdown | SFFV promoter expresses puromycin N-acetyltransferase and miR30-based shRNA targeting CAPRIN1. Target sequence: 5′-TGATTCAACTTCACTTTGCTCA-3′ | Addgene #174230 |
| pAPM miR30 CAPRIN1-ts3 | Lentivector for CAPRIN1 knockdown | SFFV promoter expresses puromycin N-acetyltransferase and miR30-based shRNA targeting CAPRIN1. Target sequence: 5′-TCTCACAAGGGGATCTGCCTGA-3′ | Addgene #174231 |
| pAPM miR30 DDX1-ts1 | Lentivector for DDX1 knockdown | SFFV promoter expresses puromycin N-acetyltransferase and miR30-based shRNA targeting DDX1. Target sequence: 5′-TAACACATGAGAACTGGTGTCC-3′ | Addgene #174232 |
| pAPM miR30 DDX1-ts2 | Lentivector for DDX1 knockdown | SFFV promoter expresses puromycin N-acetyltransferase and miR30-based shRNA targeting DDX1. Target sequence: 5′-TGGGGAGAAACTTTGTTTGTGT-3′ | Addgene #174233 |
| pAPM miR30 DDX1-ts3 | Lentivector for DDX1 knockdown | SFFV promoter expresses puromycin N-acetyltransferase and miR30-based shRNA targeting DDX1. Target sequence: 5′-TACTCTTGAACAGTAGGTGCCA-3′ | Addgene #174234 |
| pAPM miR30 DDX3X-ts1 | Lentivector for DDX3X knockdown | SFFV promoter expresses puromycin N-acetyltransferase and miR30-based shRNA targeting DDX3X. Target sequence: 5′-TAATATTTATGTTCCTCTCGTT-3′ | Addgene #174235 |
| pAPM miR30 DDX3X-ts2 | Lentivector for DDX3X knockdown | SFFV promoter expresses puromycin N-acetyltransferase and miR30-based shRNA targeting DDX3X. Target sequence: 5′-TATAGCATGCTTTTGCACTGGA-3′ | Addgene #174236 |
| pAPM miR30 DDX3X-ts3 | Lentivector for DDX3X knockdown | SFFV promoter expresses puromycin N-acetyltransferase and miR30-based shRNA targeting DDX3X. Target sequence: 5′-TAATGAGGAGGAATATAGCGCC-3′ | Addgene #174237 |

| | | | |
|---|---|---|---|
| pAPM miR30 DDX17-ts1 | Lentivector for DDX17 knockdown | SFFV promoter expresses puromycin N-acetyltransferase and miR30-based shRNA targeting DDX17. Target sequence: 5′-TAAACACAGAGAATCGGGGCTA-3′ | Addgene #174238 |
| pAPM miR30 DDX17-ts2 | Lentivector for DDX17 knockdown | SFFV promoter expresses puromycin N-acetyltransferase and miR30-based shRNA targeting DDX17. Target sequence: 5′-TGATACATCAGATTGGGATTGT-3′ | Addgene #174239 |
| pAPM miR30 DDX17-ts3 | Lentivector for DDX17 knockdown | SFFV promoter expresses puromycin N-acetyltransferase and miR30-based shRNA targeting DDX17. Target sequence: 5′-TCTGGTTGACTCTTGTCTCCAT-3′ | Addgene #174240 |
| pAPM miR30 DDX58-ts1 | Lentivector for DDX58 knockdown | SFFV promoter expresses puromycin N-acetyltransferase and miR30-based shRNA targeting DDX58. Target sequence: 5′-ATAAAGTCCAGAATAACCTGCA-3′ | Addgene #174241 |
| pAPM miR30 DDX58-ts2 | Lentivector for DDX58 knockdown | SFFV promoter expresses puromycin N-acetyltransferase and miR30-based shRNA targeting DDX58. Target sequence: 5′-TTAAATTTGTCGCTAATCCGTG-3′ | Addgene #174242 |
| pAPM miR30 DDX58-ts3 | Lentivector for DDX58 knockdown | SFFV promoter expresses puromycin N-acetyltransferase and miR30-based shRNA targeting DDX58. Target sequence: 5′-TTTCTGAACTGTAACAATCCAT-3′ | Addgene #174243 |
| pAPM miR30 DHX58-ts1 | Lentivector for DHX58 knockdown | SFFV promoter expresses puromycin N-acetyltransferase and miR30-based shRNA targeting DHX58. Target sequence: 5′-AATAGAAGTGGCCTTGGTAGGG-3′ | Addgene #174244 |
| pAPM miR30 DHX58-ts2 | Lentivector for DHX58 knockdown | SFFV promoter expresses puromycin N-acetyltransferase and miR30-based shRNA targeting DHX58. Target sequence: 5′-TTAAGTTCTAGGTACTGGCTCA-3′ | Addgene #174245 |

| | | | |
|---|---|---|---|
| pAPM miR30 DHX58-ts3 | Lentivector for DHX58 knockdown | SFFV promoter expresses puromycin N-acetyltransferase and miR30-based shRNA targeting DHX58. Target sequence: 5′-TAGACGGTGTCCTTGTGCGTGT-3′ | Addgene #174246 |
| pAPM miR30 EIF2AK2-ts1 | Lentivector for EIF2AK2 knockdown | SFFV promoter expresses puromycin N-acetyltransferase and miR30-based shRNA targeting EIF2AK2. Target sequence: 5′-TTTACTTGTTTTGTATCTACTA-3′ | Addgene #174247 |
| pAPM miR30 EIF2AK2-ts2 | Lentivector for EIF2AK2 knockdown | SFFV promoter expresses puromycin N-acetyltransferase and miR30-based shRNA targeting EIF2AK2. Target sequence: 5′-TTAACTTGAAATGTAAACCTCC-3′ | Addgene #174248 |
| pAPM miR30 EIF2AK2-ts3 | Lentivector for EIF2AK2 knockdown | SFFV promoter expresses puromycin N-acetyltransferase and miR30-based shRNA targeting EIF2AK2. Target sequence: 5′-TCTATTTTTGCTGTTCTCAGGA-3′ | Addgene #174249 |
| pAPM miR30 HNRNPK-ts1 | Lentivector for HNRNPK knockdown | SFFV promoter expresses puromycin N-acetyltransferase and miR30-based shRNA targeting HNRNPK. Target sequence: 5′-TAATTCAACCATCTCATCAGTG-3′ | Addgene #174250 |
| pAPM miR30 HNRNPK-ts2 | Lentivector for HNRNPK knockdown | SFFV promoter expresses puromycin N-acetyltransferase and miR30-based shRNA targeting HNRNPK. Target sequence: 5′-TTAATCATAGGTTTCATCGTAA-3′ | Addgene #174251 |
| pAPM miR30 HNRNPK-ts3 | Lentivector for HNRNPK knockdown | SFFV promoter expresses puromycin N-acetyltransferase and miR30-based shRNA targeting HNRNPK. Target sequence: 5′-TAATCTTTGGGAATAGTTACTT-3′ | Addgene #174252 |

| | | | |
|---|---|---|---|
| pAPM miR30 HNRNPR-ts1 | Lentivector for HNRNPR knockdown | SFFV promoter expresses puromycin N-acetyltransferase and miR30-based shRNA targeting HNRNPR. Target sequence: 5′-TTACATCCAGAGTATAACCAGT-3′ | Addgene #174253 |
| pAPM miR30 HNRNPR-ts2 | Lentivector for HNRNPR knockdown | SFFV promoter expresses puromycin N-acetyltransferase and miR30-based shRNA targeting HNRNPR. Target sequence: 5′-TTTCATAGCTGTCACACAGTTT-3′ | Addgene #174254 |
| pAPM miR30 HNRNPR-ts3 | Lentivector for HNRNPR knockdown | SFFV promoter expresses puromycin N-acetyltransferase and miR30-based shRNA targeting HNRNPR. Target sequence: 5′-TACAAAAAGTCTGTTGTTTGCC-3′ | Addgene #174255 |
| pAPM miR30 IFI16-ts1 | Lentivector for IFI16 knockdown | SFFV promoter expresses puromycin N-acetyltransferase and miR30-based shRNA targeting IFI16. Target sequence: 5′-TTGAATTTCTCCTTCAAGCTGG-3′ | Addgene #174256 |
| pAPM miR30 IFI16-ts2 | Lentivector for IFI16 knockdown | SFFV promoter expresses puromycin N-acetyltransferase and miR30-based shRNA targeting IFI16. Target sequence: 5′-TTTAGTAGAACAATGTTCTTGT-3′ | Addgene #174257 |
| pAPM miR30 IFI16-ts3 | Lentivector for IFI16 knockdown | SFFV promoter expresses puromycin N-acetyltransferase and miR30-based shRNA targeting IFI16. Target sequence: 5′-TAGTAGAACAATGTTCTTGTAT-3′ | Addgene #174258 |
| pAPM miR30 IFIH1-ts1 | Lentivector for IFIH1 knockdown | SFFV promoter expresses puromycin N-acetyltransferase and miR30-based shRNA targeting IFIH1. Target sequence: 5′-TTGTGTCATTAATTTGTAGGGC-3′ | Addgene #174259 |
| pAPM miR30 IFIH1-ts2 | Lentivector for IFIH1 knockdown | SFFV promoter expresses puromycin N-acetyltransferase and miR30-based shRNA targeting IFIH1. Target sequence: 5′-TTTATACATCATCTTCTCTCGG-3′ | Addgene #174260 |

| | | | |
|---|---|---|---|
| pAPM miR30 IFIH1-ts3 | Lentivector for IFIH1 knockdown | SFFV promoter expresses puromycin N-acetyltransferase and miR30-based shRNA targeting IFIH1. Target sequence: 5′-ATGTTACATTCTTTAATATCCA-3′ | Addgene #174261 |
| pAPM miR30 ILF2-ts1 | Lentivector for ILF2 knockdown | SFFV promoter expresses puromycin N-acetyltransferase and miR30-based shRNA targeting ILF2. Target sequence: 5′-TAAAACTTCAGAAGGATCCTGT-3′ | Addgene #174262 |
| pAPM miR30 ILF2-ts2 | Lentivector for ILF2 knockdown | SFFV promoter expresses puromycin N-acetyltransferase and miR30-based shRNA targeting ILF2. Target sequence: 5′-TTGTAATGAGAATCTTCACTGT-3′ | Addgene #174263 |
| pAPM miR30 ILF2-ts3 | Lentivector for ILF2 knockdown | SFFV promoter expresses puromycin N-acetyltransferase and miR30-based shRNA targeting ILF2. Target sequence: 5′-TGAACTTTAACTGTGGACTGAG-3′ | Addgene #174264 |
| pAPM miR30 ILF3-ts1 | Lentivector for ILF3 knockdown | SFFV promoter expresses puromycin N-acetyltransferase and miR30-based shRNA targeting ILF3. Target sequence: 5′-TTTTGTTGGAACCAGCACCTTG-3′ | Addgene #174265 |
| pAPM miR30 ILF3-ts2 | Lentivector for ILF3 knockdown | SFFV promoter expresses puromycin N-acetyltransferase and miR30-based shRNA targeting ILF3. Target sequence: 5′-TATTTCGTACTTGTCTTCTGTT-3′ | Addgene #174266 |
| pAPM miR30 ILF3-ts3 | Lentivector for ILF3 knockdown | SFFV promoter expresses puromycin N-acetyltransferase and miR30-based shRNA targeting ILF3. Target sequence: 5′-TTCTTTACTGTCGTCCTCTGGG-3′ | Addgene #174267 |
| pAPM miR30 LRPPRC-ts1 | Lentivector for LRPPRC knockdown | SFFV promoter expresses puromycin N-acetyltransferase and miR30-based shRNA targeting LRPPRC. Target sequence: 5′-TGTAATTTAATGAGATTGCTGT-3′ | Addgene #174268 |

| | | | |
|---|---|---|---|
| pAPM miR30 LRPPRC-ts2 | Lentivector for LRPPRC knockdown | SFFV promoter expresses puromycin N-acetyltransferase and miR30-based shRNA targeting LRPPRC. Target sequence: 5′-TATCCAAGAATCTTGCTGGCAC-3′ | Addgene #174269 |
| pAPM miR30 LRPPRC-ts3 | Lentivector for LRPPRC knockdown | SFFV promoter expresses puromycin N-acetyltransferase and miR30-based shRNA targeting LRPPRC. Target sequence: 5′-TTTATATTCATAGACCTCCTGA-3′ | Addgene #174270 |
| pAPM miR30 MATR3-ts1 | Lentivector for MATR3 knockdown | SFFV promoter expresses puromycin N-acetyltransferase and miR30-based shRNA targeting MATR3. Target sequence: 5′-TTCGGTTGAACTCTCAGTCTTC-3′ | Addgene #174271 |
| pAPM miR30 MATR3-ts2 | Lentivector for MATR3 knockdown | SFFV promoter expresses puromycin N-acetyltransferase and miR30-based shRNA targeting MATR3. Target sequence: 5′-TAATTAGCAAGATCGGTCTCGT-3′ | Addgene #174272 |
| pAPM miR30 MATR3-ts3 | Lentivector for MATR3 knockdown | SFFV promoter expresses puromycin N-acetyltransferase and miR30-based shRNA targeting MATR3. Target sequence: 5′-TTAGCAAGATCGGTCTCGTCTC-3′ | Addgene #174273 |
| pAPM miR30 MAVS-ts1 | Lentivector for MAVS knockdown | SFFV promoter expresses puromycin N-acetyltransferase and miR30-based shRNA targeting MAVS. Target sequence: 5′-TTAGAAGGCACTGCACCAGGGG-3′ | Addgene #174274 |
| pAPM miR30 MAVS-ts2 | Lentivector for MAVS knockdown | SFFV promoter expresses puromycin N-acetyltransferase and miR30-based shRNA targeting MAVS. Target sequence: 5′-TTTGGATGGTGCTGGATTGGTG-3′ | Addgene #174275 |

| | | | |
|---|---|---|---|
| pAPM miR30 MAVS-ts3 | Lentivector for MAVS knockdown | SFFV promoter expresses puromycin N-acetyltransferase and miR30-based shRNA targeting MAVS. Target sequence: 5′-TAACTTGGCTCCTTCTCTCTGC-3′ | Addgene #174276 |
| pAPM miR30 MOV10-ts1 | Lentivector for MOV10 knockdown | SFFV promoter expresses puromycin N-acetyltransferase and miR30-based shRNA targeting MOV10. Target sequence: 5′-TATTAAGACCCGGTATTCCTGC-3′ | Addgene #174277 |
| pAPM miR30 MOV10-ts2 | Lentivector for MOV10 knockdown | SFFV promoter expresses puromycin N-acetyltransferase and miR30-based shRNA targeting MOV10. Target sequence: 5′-TAACGTGACAGTCTTGCCGGTG-3′ | Addgene #174278 |
| pAPM miR30 MOV10-ts3 | Lentivector for MOV10 knockdown | SFFV promoter expresses puromycin N-acetyltransferase and miR30-based shRNA targeting MOV10. Target sequence: 5′-TAAGCTGATCTTGAAGTCGCGG-3′ | Addgene #174279 |
| pAPM miR30 MYD88-ts1 | Lentivector for MYD88 knockdown | SFFV promoter expresses puromycin N-acetyltransferase and miR30-based shRNA targeting MYD88. Target sequence: 5′-TTCTTCATTGCCTTGTACTTGA-3′ | Addgene #174280 |
| pAPM miR30 MYD88-ts2 | Lentivector for MYD88 knockdown | SFFV promoter expresses puromycin N-acetyltransferase and miR30-based shRNA targeting MYD88. Target sequence: 5′-TTGGTGTAGTCGCAGACAGTGA-3′ | Addgene #174281 |
| pAPM miR30 MYD88-ts3 | Lentivector for MYD88 knockdown | SFFV promoter expresses puromycin N-acetyltransferase and miR30-based shRNA targeting MYD88. Target sequence: 5′-TCAGTCGATAGTTTGTCTGTTC-3′ | Addgene #174282 |

| | | | |
|---|---|---|---|
| pAPM miR30 TMEM173-ts1 | Lentivector for TMEM173 knockdown | SFFV promoter expresses puromycin N-acetyltransferase and miR30-based shRNA targeting TMEM173. Target sequence: 5′-TACAAAGTCTGCAAGGGGGTGG-3′ | Addgene #174283 |
| pAPM miR30 TMEM173-ts2 | Lentivector for TMEM173 knockdown | SFFV promoter expresses puromycin N-acetyltransferase and miR30-based shRNA targeting TMEM173. Target sequence: 5′-TAAGCCAGCTTGACTGTATTGT-3′ | Addgene #174284 |
| pAPM miR30 TMEM173-ts3 | Lentivector for TMEM173 knockdown | SFFV promoter expresses puromycin N-acetyltransferase and miR30-based shRNA targeting TMEM173. Target sequence: 5′-TAATGCTGATTGTAAGTTCGAA-3′ | Addgene #174285 |
| pAPM miR30 XPO1-ts1 | Lentivector for XPO1 knockdown | SFFV promoter expresses puromycin N-acetyltransferase and miR30-based shRNA targeting XPO1. Target sequence: 5′-TTCTGTATCTACATAATCCAGA-3′ | Addgene #174286 |
| pAPM miR30 XPO1-ts2 | Lentivector for XPO1 knockdown | SFFV promoter expresses puromycin N-acetyltransferase and miR30-based shRNA targeting XPO1. Target sequence: 5′-TATAAGTTGATCATGTTCCTTA-3′ | Addgene #174287 |
| pAPM miR30 XPO1-ts3 | Lentivector for XPO1 knockdown | SFFV promoter expresses puromycin N-acetyltransferase and miR30-based shRNA targeting XPO1. Target sequence: 5′-TAACAGTAACAAGAAATCGTTT-3′ | Addgene #174288 |
| pAPM miR30 ZC3HAV1-ts1 | Lentivector for ZC3HAV1 knockdown | SFFV promoter expresses puromycin N-acetyltransferase and miR30-based shRNA targeting ZC3HAV1. Target sequence: 5′-ATAACTTTTGCATATCTCGGGC-3′ | Addgene #174289 |

| | | | |
|---|---|---|---|
| pAPM miR30 ZC3HAV1-ts2 | Lentivector for ZC3HAV1 knockdown | SFFV promoter expresses puromycin N-acetyltransferase and miR30-based shRNA targeting ZC3HAV1. Target sequence: 5′-TTAACAAGTGCTACTCTTGGGT-3′ | Addgene #174290 |
| pAPM miR30 ZC3HAV1-ts3 | Lentivector for ZC3HAV1 knockdown | SFFV promoter expresses puromycin N-acetyltransferase and miR30-based shRNA targeting ZC3HAV1. Target sequence: 5′-TTGATGATCCTGTAGTCTGAGG-3′ | Addgene #174291 |

**Supplementary Table 4. Oligonucleotides used in RT-qPCR of HIV-1 following formaldehyde cross-linking immunoprecipitation (fCLIP)**

| Name | Sequence |
|---|---|
| 5'LTR-F | 5'-TGTTCGGGCGCCACTGCTAGAGA-3' |
| 5'LTR-R | 5'-GGAACCCACTGCTTAAGCCTCAA-3' |
| gag-F | 5'-ACATCAAGCAGCCATGCAAAT-3' |
| gag-R | 5'-TCTGGCCTGGTGCAATAGG-3' |
| RT-F | 5'-AGAAATACAGAAGCAGGGGCA-3' |
| RT-R | 5'-TGGGCACCCTTCATTCTTGC-3' |
| IN-F | 5'-TGGAGAGCAATGGCTAGTGA-3' |
| IN-R | 5'-GCATGGCTTCCCCTTTTAGC-3' |
| D1A4-F | 5'-GATCTCTCGACGCAGGACTC-3' |
| D1A4-R | 5'-TGGTCCTTTCCAAACTGGAT-3' |
| D4A7-F | 5'-CAGGAAGAAGCGGAGACAG-3' |
| D4A7-R | 5'-ATTGGGAGGTGGGTCTGAAAC-3' |
| ACTB-F | 5'-GTGATGGTGGGCATGGGTC-3' |
| ACTB-R | 5'-CCATGTCGTCCCAGTTGGTG-3' |
